# Destabilized 3’UTR ARE therapeutically degrades ERBB2 in drug-resistant ERBB2+ cancer models

**DOI:** 10.1101/2022.08.14.503914

**Authors:** Chidiebere U Awah, Yana Glemaud, Fayola Levine, Kiseok Yang, Afrin Ansary, Fu Dong, Leonard Ash, Junfei Zhang, Daniel Weiser, Olorunseun O Ogunwobi

## Abstract

Breast, lung, and colorectal cancer resistance to molecular targeted therapy is a major challenge and unfavorably impacts clinical outcomes, leading to hundreds of thousands of deaths yearly. In ERBB2+ cancers regardless of the tissue of origin, ERBB2 is the driver oncogene of resistance. We discovered that the ERBB2+ cancers are enriched with poly U sequences on their 3’UTR AU rich elements which are mRNA stabilizing sequences. We developed a novel technology, in which we engineered these ERBB2 mRNA stabilizing sequences to unstable forms and specifically controlled and degraded ERBB2 transcript and protein across multiple cancer types both in the wildtype and drug resistance settings *in vitro* and *in vivo*, offering a unique novel modality to control ERBB2 and other pervasive oncogenic signals where other therapies fail.

**One-Sentence Summary:** Engineered destabilized 3’UTR ARE of ERBB2 degrades ERBB2 in many cancer types and controlled resistance.

**Graphical abstract:** **A.** Depiction represents multiple ERBB2 expressing cancer cells with stable 3’UTR ARE and the signaling cascade known to cause chemo resistance**. B.** Depiction of the engineered destabilized 3’UTR ARE of ERBB2 and the destabilization and degradation of the ERBB2 transcript, protein and kinases involved in mediation of drug resistance

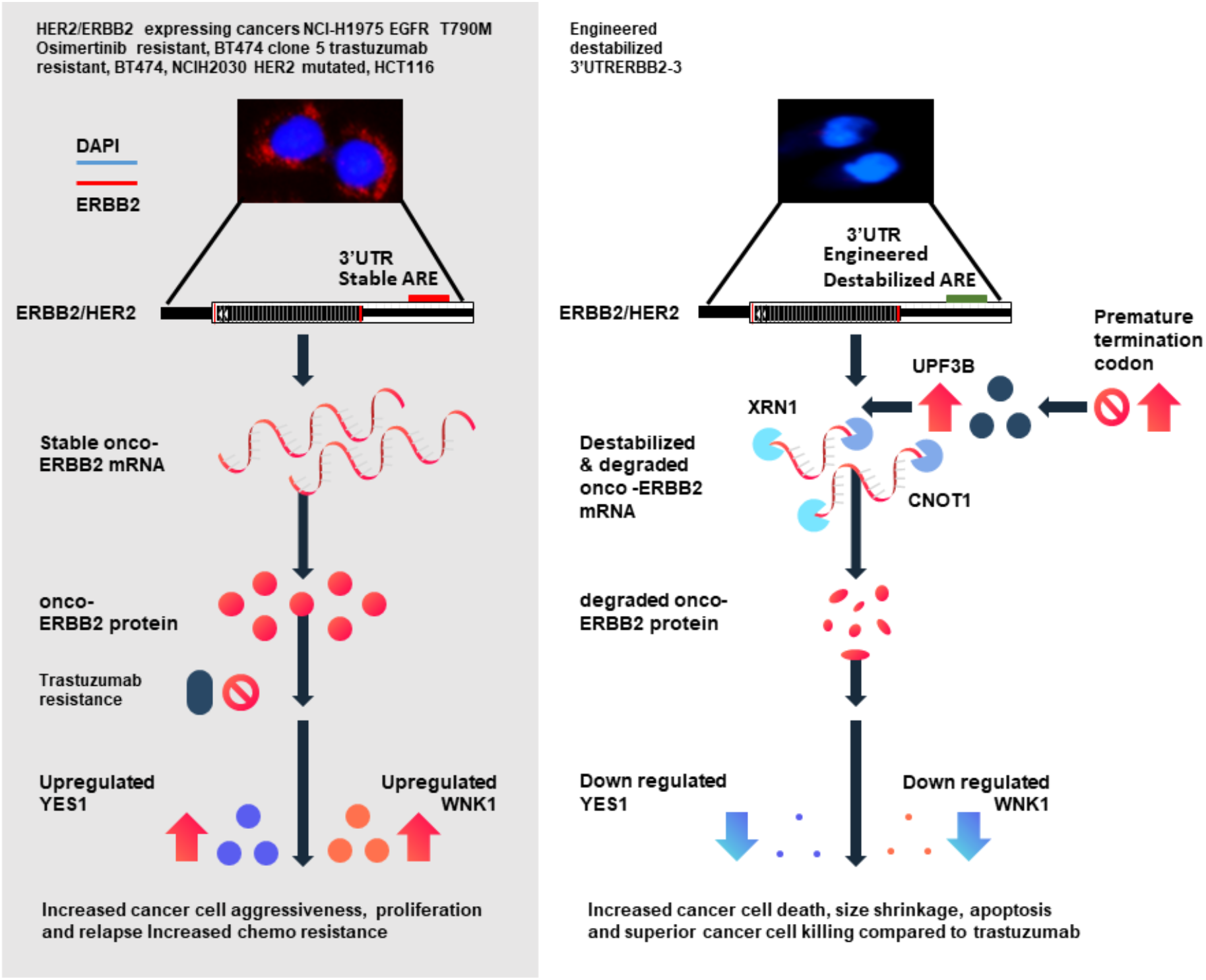

---

Breast and lung cancers are deadly tumors afflicting millions of patients worldwide annually. Breast cancer is the most common form of cancer in women worldwide with mortality of about 58% in developing countries (***1***). According to NCI SEER statistics, 53.1 per 100,000 men and women will develop lung and bronchial cancer per year, with 5-year survival rate at 21.7%. Lung cancer caused more deaths in 2017 than breast cancer, prostate cancer and brain cancers combined (***2***). In the US, the incidence of breast cancer is at 1.6% annual growth rate (AGR) at 267,866 cases per year. In Western Europe (Germany, France, UK), the incidence of breast cancer is between 0.39-0.98% averaging about 50,000 to 77,000 cases. In China breast cancer is at 3.36% with 244,949 cases per year while incidence rates in Japan at 90,452 cases per year. In Africa, breast cancer is at 28% incidence with very poor outcomes and high mortality accounting for 20% of all cancer deaths in women (***3***). There is disparity in breast cancer survival amongst women of African descent when compared to women of other ancestry. A recent study shows that there is significant disparity across all survival outcome indices in women of African ancestry with ERBB2+ breast cancer in comparison to women of other ancestry with ERBB2+ breast cancer (***4***).The prevalence of breast and lung cancer and their related mortality arises from the oncogenes that drive the malignant breast and lung tissues to excessively proliferate and then metastasize to lymph nodes and onwards to organs such as lungs, liver, and brain leading to death of the patient.

ERBB2/ERBB2 is a member of the subclass I receptor tyrosine kinase superfamily of ERBB/ EGFR (epidermal growth factor receptor family) which consist of four members namely: EGFR/ERBB1, ERBB2, ERBB3 and ERBB4 (***5, 6***). ERBB2 is activated upon binding of the neuregulin ligand unto the ERBB receptor which leads to its homo and heterodimerization triggering the activation of tyrosine kinases which have docking sites on the ERBB receptors and from there control a large-scale signaling proteins, transcription factors and kinases which mediates ERBB functions. ERBB2/ERBB2 is overexpressed in many human cancers including the breast, lung and colorectal cancers. The overexpression of ERBB2 is associated with very aggressive breast and drug resistant lung cancer because ERBB2 is a membrane protein that signals and amplifies for proliferation, pro-survival and prometastatic signals of the cancer leading to poor clinical outcomes.

Tyrosine kinase inhibitors such as osimertinib, erlotinib and gefitinib target the ERRB/EGFR and antibodies such as trastuzumab which target the ecto-domain of ERBB2 have been developed as therapeutics such as ERBB2 overexpressing cancers. Trastuzumab in various forms in combination with chemotherapy has been successful in targeting ERBB2 overexpression in ERBB2+ breast cancer and improving survival as a standard first line therapy for more than a decade (***7-8***). However, there are groups of ERBB2+ breast cancer, lung cancer, esophageal cancers and gastrointestinal cancers that are resistant to trastuzumab and tyrosine kinase inhibitors (***9-21***). It is well known that 25% of early stage and 75% of late-stage ERBB2+ breast cancer is resistant to trastuzumab. Same is true for EGFR T790M non-small cell lung cancer that are osimertinib resistant with 6% of patients bearing ERBB2 amplification.

For these patients who have treatment resistant cancers with overexpression of ERBB2, there is limited clinical intervention for them. Moreso, for these trastuzumab resistant ERBB2 driven cancer patients, regional and distant organ metastasis of the tumor is a significant obstacle to good clinical outcome. Resistance to trastuzumab and other tyrosine kinase inhibitors in ERBB2+ breast and lung cancer is a grand challenge that needs to be overcome to improve clinical outcomes, and to prevent drug resistance and help sufferers of these deadly cancers.

To address this grand challenge of ERBB2+ drug resistance in breast and lung cancer, we have developed an innovative approach based on genome engineering of the genetic codes on the 3’UTR (3’ untranslated region) of the ERBB2 gene to destabilize and degrade ERBB2 transcript, protein expression, ERBB2 dependent kinases and interactome. The structure of a typical gene consists of various parts namely 5’UTR (untranslated region), the CDS (coding sequence), and the end of the gene called 3’UTR. The 3’UTR of genes serves as a hub for translation, stabilizing and destabilizing of transcript and polyadenylation (***22***). Various motifs have been discovered in the 3’UTR region of genes to determine which transcripts are stabilized or destabilized and degraded (***23, 24***). The AU rich elements (ARE) found on the 3’UTR have been demonstrated to be involved in nonsense mediated decay of transcript via a complex process that involves decapping enzymes, deadenylation proteins and cleaving enzymes (***25***). Proteins such as the human DCP1A, DCP2 and ZFP36 are involved in this process of ARE mediated decay (***25***). Motifs on the 3’UTR of the gene were found to control mRNA transcript stabilization or destabilization. (***23-24***). The consensus stabilization motifs are UUUUU, UUGCAUGG, and CCUUACAC whereas the destabilization motifs are AUUUU, CCUC, CUGC, UAAGUUAU, UAACUUAU and GUAAAUAG.

Based on these findings, we hypothesized that the 3’UTR of oncogenes such as ERBB2 will be enriched with ARE stabilizing elements which stabilizes their oncogenic transcript and drives tumor aggressiveness. Therefore, if we change these stabilizing elements to destabilizing elements by motif engineering and driven by mRNA decapping promoter DCP1A that we can control the oncogene by degrading the oncogenic transcript, proteins and signaling cascade. Here, we provide proof of principle results that by destabilizing the stable 3’UTR elements of ERBB2 across multiple recalcitrant drug resistant cancer types, that we controlled pervasive ERBB2 oncogenic transcript and specifically degraded the ERBB2 protein and its associated kinases and interactome which triggered apoptosis and killed the tumors and is validated *in vivo*. These findings uncover a novel technology to target and control oncogenes where drugs failed and for therapeutic gene targeting in diseases and control of any transcript of interest.

## Results

### Identification of poly U stabilizing ARE motifs and design of the engineered destabilized 3’UTR ARE ERBB2 constructs

We hypothesized that the oncogenic transcript of ERBB2 is stabilized on the 3’UTR with poly U sequence and that if the stabilizing element is destabilized, we could degrade the ERBB2 transcript and protein expression, and thus control the aggressiveness of ERBB2+ driven cancers both in wildtype and drug resistant settings. To do this, (**Fig. 1A**), we extracted RNA from ERBB2+ and non-ERBB2+ breast cancer cells (BT474, MCF7, MDA MB231 and T47D), performed cDNA synthesis, performed PCR amplification of the cDNA, and then sequenced them with ERBB2 3’UTR primers by Sanger sequencing (**Fig. S3A-E**). We found that the 3’UTR of ERBB2 in only ERBB2 expressing cancer cells MCF7, BT474 and T47D (**Fig. S3A, B, C and E**) are enriched with UUUUUU sequences which are stabilizing AU rich elements (**Fig. 1A, Fig. S3A, B, C, E and Fig. S4A**). But these stabilizing elements are not enriched in triple negative breast cancer MDA MB231 (**Fig. S3D**). We designed a synthetic g-block of this 3’UTR region wherein we engineered the various stabilizing elements into destabilized elements and cloned them into a vector (**Fig. 1A, Fig. S4A, Fig. S5A-B and Fig. S6A-B**). To design the destabilized 3’UTR ARE, we replaced all the stable elements with the destabilized ARE elements which are CCUC, CUGC, UAAGUUAU (**Fig. S4A**). We reasoned that if we destabilized the ERBB2-3’UTR ARE and drive its expression by the deadenylating enzyme promoter DCP1A, that we could degrade the transcript and protein specifically through nonsense mediated decay. Therefore, we designed the destabilizing ERBB2 synthetic g block to contain BstB1 site, a poly A sequence to stop RFP transcription and then a DCP1A promoter element to drive the destabilized ERBB2 3’UTR ARE and then on the 3’ end with Bamh1 site and a poly A to stop DCP1A (**Fig. S6A-B**).

**Figure 1:**
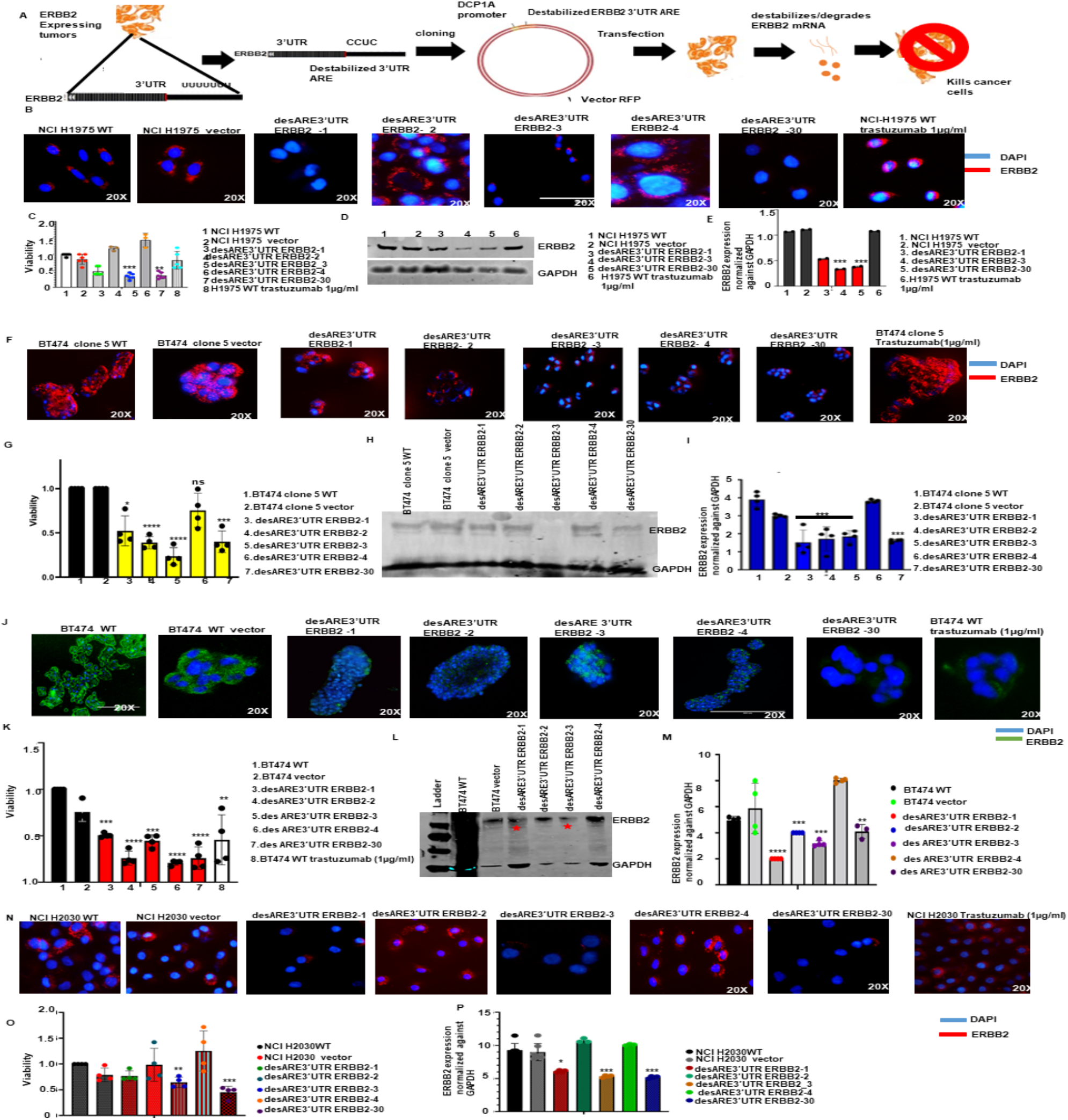
The destabilized 3’UTR ARE of ERBB2 degrades oncogenic ERBB2 protein and transcript in EGFR T90M mutated lung cancer, in ERBB2+ trastuzumab resistant breast cancer, in wild type ERBB2+ breast cancer and in ERBB2 mutated lung cancer leading to cell death. **A.** Schematic depiction of the destabilization technology, ERBB2 expressing cancers RNA is extracted and 3’UTR cDNA is made and sequenced by Sanger method, the poly U sequences are identified. The poly U sequences on the 3’UTR are engineered and changed to the destabilized motif which is cloned into an RFP vector and driven by DCP1A. The constructs are transfected into ERBB2 expressing cells and mRNA transcript destabilization is established within 4 days, the cellular membranes denude, and the cancer cell is killed within 8 days. **B.** Immunofluorescence pictures show ERBB2 expression (stained in red) and nuclei (DAPI-blue) on wildtype NCI-H1975, vector and on NCI-H1975 containing constructs desARE3’UTR ERBB2-1, 2, 3, 4, 30 and wild type treated trastuzumab NCI-H1975 (n=2). **C.** The bar charts show cells viability of the NCI-H1975 wildtype cells, vector and cells containing constructs desARE3’UTR ERBB2-1, 2, 3,4 and 30 and wildtype trastuzumab treated cells (n=2). Paired two tailed T-test (***Pvalue=0.0013 desARE3’UTRERBB2-3, **Pvalue= 0.03 desARE3’UTRERBB2-30). **D.** Western blot show ERBB2 and GAPDH expression across the NCI-H1975 wildtype cells, vector, cells containing constructs desARE3’UTR ERBB2-1, 3 and 30 and wild type trastuzumab treated NCI-H1975 cells (n=2). **E.** Bar charts show quantification of the ERBB2 expression normalized against GAPDH by qPCR on NCI-H1975 wildtype cells, vector and cells containing constructs desARE3’UTR ERBB2-1, 2, 3, 4, and 30 and wild type trastuzumab treated NCI-H1975 cells (n=2). Paired two tailed T-test (***Pvalue=0.007569 desARE3’UTRERBB2-3, ***Pvalue=0.001123 desARE3’UTR ERBB2-30). **F.** Immunofluorescence images show ERBB2 expression (stained in red) and nuclei (DAPI-blue) on wildtype BT474 clone 5, vector and on BT474 clone 5 containing constructs desARE3’UTR ERBB2-1, 2, 3, 4, 30 and BT474 clone 5 WT trastuzumab treated (n=2). **G.** The bar charts show cells viability of the BT474 clone 5 wildtype cells, vector and cells containing constructs desARE3’UTR ERBB2-1, 2, 3, 4 and 30 (n=2). Paired two tailed T-test (**Pvalue=0.011034 desARE3’UTRERBB2-1, ****Pvalue=0.000427 desARE3’UTRERBB2-2, ****Pvalue=0.000579 desARE3’UTRERBB2-3, ns=0.084504 desARE3’UTRERBB2-4, ***Pvalue=0.002188 desARE3’UTRERBB2-30). **H.** Western blot show ERBB2 and GAPDH expression across the wildtype BT474 clone 5 WT, vector and on BT474 clone 5 containing constructs desARE3’UTR ERBB2-1, 2, 3 and 4 (n=3). **I.** Bar charts show quantification of the ERBB2 expression normalized against GAPDH by qPCR on BT474 clone 5 wildtype cells, vector and BT474 clone 5 cells containing constructs desARE3’UTR ERBB2-1, 2, 3, 4 and 30 (n=2). Paired two tailed T-test (***=0.003, desARE3’UTRERBB2-1, 2, 3 and 30). **J.** Immunofluorescence images show ERBB2 expression (stained in green) and nuclei (DAPI-blue) on wildtype BT474, vector and on BT474 containing constructs desARE3’UTR ERBB2-1, 2, 3, 4, 30 and BT474 wild type trastuzumab treated cells (n=2). **K.** The bar charts show cells viability of the BT474 wildtype cells, vector and BT474 cells containing constructs desARE3’UTR ERBB2-1, 2, 3, 4 (n=4). Paired two tailed T-test (***Pvalue=0.0025desARE3’UTRERBB2-1,****Pvalue=0.00029515 desARE3’UTRERBB2-2,***Pvalue=0.0045desARE3’UTRERBB2-3, ****Pvalue=0.000102desARE3’UTRERBB2-4,***Pvalue=0.00120 desARE3’UTRERBB2-30, **Pvalue=0.027 BT474 WT trastuzumab treated. **L.** Western blots show ERBB2 and GAPDH expression across the wildtype BT474, vector and on BT474 containing constructs desARE3’UTR ERBB2-1, 2, 3 and 4 (n=2). **M.** Bar charts show quantification of the ERBB2 expression normalized against GAPDH by qPCR on BT474 wildtype cells, vector and cells containing constructs desARE3’UTR ERBB2-1, 2, 3, and 30 (n=2). Paired two tailed T-test (****P value=0.0003 desARE3’UTRERBB2-1, ***Pvalue=0.0021 desARE3’UTRERBB2-2,3, **P value=0.013 desARE3’UTRERBB2-30). **N.** Immunofluorescence images show ERBB2 expression (stained in red) and nuclei (DAPI-blue) on wildtype NCI H2030 cells, vector and on NCI-H2030 containing constructs desARE3’UTR ERBB2-1, 2, 3, 4, 30 and trastuzumab treated cells (n=2). **O.** The bar charts show cells viability of the NCI H2030 wildtype cells, vector and cells containing constructs desARE3’UTR ERBB2-1,2,3, 4 and 30 (n=2). Paired two tailed T-test (** Pvalue =0.0080 desARE3’UTRERBB2-3, *** =0.0024 desARE3’UTRERBB2-30). **P.** Bar charts show quantification of the ERBB2 expression normalized against GAPDH by qPCR on NCI H2030 wildtype cells, vector and cells containing constructs desARE3’UTR ERBB2-1, 2, 3, 4 and 30 (n=2). Paired two tailed T-test (* Pval=0.027 desARE3’UTRERBB2-1, *** P value =0.0003 desARE3’UTRERBB2-3, *** P value =0.0001 desARE3’UTRERBB2-30.

### Engineered destabilized 3’UTR ARE of ERBB2 degrades ERBB2 in ERBB2 expressing EGFR T790M lung cancer cells, ERBB2+ trastuzumab resistant breast cancer cells, in wild type ERBB2+ breast cancer cells, in ERBB2 mutated lung cancer cells, and in ERBB2 expressing colorectal cancer cells

Our cloning strategy yielded four destabilizing ARE 3’UTR ERBB2 clones by sticky end cloning in recombinant competent E. coli namely: desARE3’UTR ERBB2-1, desARE3’UTR ERBB2-2, desARE 3’UTR ERBB2-3 and desARE3’UTR ERBB2-4 (**Fig S7A-D**) and a clone by Gibson assembly in recombinant deficient E. coli: desARE3’UTR ERBB2-30 (**Fig. S8A**). We transfected the four destabilized (des) clones (desARE3’UTRERBB2-1, desARE3’UTR ERBB2-2, desARE3’UTR ERBB2-3, desARE3’UTRERBB2-4 and desARE 3’UTR ERBB2-30 into non-small cell lung cancer cells that carry mutation in the EGFR T790M but expresses ERBB2. Within 4 days (**Fig. 1B**), by immunofluorescence staining for ERBB2 (ERBB2 stained in red and nuclei stained blue with DAPI) and western blotting detection of ERBB2 (**Fig. 1D**), we found that ERBB2 protein expression was lost in desARE3’UTR ERBB2-1, 3 and 30 compared to the wild type cells, vector control and trastuzumab treated NCI H1975 cells. The NCI H1975 cells transfected with desARE3’UTR ERBB2-1, 3 and 30 show significant loss of viability compared to the controls (**Fig. 1C**). In western blot the desARE3’UTR ERBB2-3 and 30 (**Fig. 1D**) caused loss of ERBB2 protein expression compared to controls. The desARE3’UTR ERBB2-1, 3 and 30 also caused significant decrease in ERBB2 transcript expression compared to controls (**Fig. 1E**). The morphology of the destabilized cells shows ruptured membrane compared to controls (**Fig. S1A**). The loss of ERBB2 at the protein level as determined by immunofluorescence and western blot were quantified and displayed in (**Fig. S1B**). We found the increased expression of cleaved caspase 3 and 9 in the desARE3’UTRERBB2-1, 3, 30 treated cells (**Fig. S2D, E, F, G, H**), and treated cells show significant inhibition of migration as demonstrated by significantly impaired wound healing (**Fig. S2I, J**). Taken together, these results demonstrate that the engineered destabilized 3’UTR ARE of ERBB2 targeted, destabilized and degraded ERBB2 transcript and protein expression, and induced increased apoptosis and reduced migration and it outperformed trastuzumab in the NCI H1975 model of the deadly EGFRT790M non-small cell lung cancer.

Having established that the ERBB2 destabilizing constructs work and degrades ERBB2 transcript and protein expression, we turned our attention to the devastating clinical problem of ERBB2+ trastuzumab resistant cancers. Trastuzumab has improved overall survival outcomes for ERBB2+ breast cancer. However, more than 25% of early stage ERBB2+ breast cancer patients develop resistance to trastuzumab and more than 75% late stage ERBB2+ breast cancers are resistant to trastuzumab even if given in combination with anthracyclines. Moreso, there is no effective treatment for drug resistance in ERBB2+ breast cancer tumor relapse. To ascertain if the novel engineered destabilized 3’UTR ARE of ERBB2 will successfully target ERBB2 in trastuzumab resistant breast cancer cells and control its aggressiveness, we experimented with the BT474 clone 5 (**26-27**) ERBB2+ trastuzumab resistant breast cancer cell line developed by prolonged exposure to trastuzumab. We transfected these cells with the ERBB2 3’UTR destabilizing constructs and within 9 days (**Fig. 1F**), we found by immunofluorescence analysis that ERBB2 (stained in red) in the treated cells is diminished compared to the wildtype cells and vector controls (**Fig. 1F**, **Fig. S1E**). We confirmed this by western blotting (**Fig. 1H**) and showed that desARE3’UTR ERBB2-3 significantly degraded ERBB2 expression and desARE3’UTR ERBB2-30 led to more than 70% loss of ERBB2 protein expression in the ERBB2+ trastuzumab resistant breast cancer cells compared to wildtype and vector control cells (**Fig. 1H, Fig. S1F**). The treated cells (except for those treated with desARE3’UTR ERBB2-4) also showed loss of viability compared to the controls (**Fig. 1G**). The morphology of the trastuzumab resistant ERBB2+ cancer cells containing the destabilized elements show shrinking and reduction of cell size compared to the controls (**Fig. 1F**, **Fig. S1D**). Transcript analysis by quantitative reverse transcriptase polymerase chain reaction (qPCR) showed that all the ERBB2 3’UTR destabilizing constructs (**Fig. 1I**) significantly reduced ERBB2 transcript, except for desARE3’UTR ERBB2-4 compared to the controls. ERBB2 3’UTR destabilizing constructs induced increased caspase 3 in the trastuzumab resistant ERBB2+ cancer cells (**Fig. S1G**). Again, this result demonstrates that the engineered destabilizing ERBB2 3’UTR ARE outperformed trastuzumab in the drug resistant setting.

To further validate this, we transfected the four clones (desARE3’UTR ERBB2-1, desARE3’UTR ERBB2-2, desARE 3’UTR ERBB2-3 and desARE3’UTR ERBB2-4) of the engineered destabilized 3’UTR ARE of ERBB2 into ERBB2+ BT474 cells. Within 4 days (**Fig. 1J, Fig. S1I**), we observed by immunofluorescence that ERBB2 protein (stained green) expression has been significantly decreased in the cancer cells containing the destabilizing elements compared to the wildtype and vector control. The morphology of the destabilized cells under microscope shows distorted and ruptured membrane compared to the wildtype cells and the vector control (**Fig. S1H**). To confirm the loss of ERBB2 protein, we performed western blotting on the wildtype, vector controls cells and the treated cells (**Fig. 1L, Fig. S1J**). We found that the desARE3’UTR ERBB2-1 and desARE3’UTR ERBB2-3 degraded ERBB2 protein (**Fig. 1L, Fig. S1J**) and transcript (**Fig. 1M**) expression. Within 8 days, all the cancer cells transfected with ERBB2 3’UTR destabilizing elements showed loss of viability and increased caspase 3/7 expression (**Fig. S1K**) compared to the wildtype cells and the vector control (**Fig. 1K**).

To further demonstrate the generalizability of the novel engineered 3’UTR ARE destabilization of ERBB2 oncogene, we extended this work into the NCIH2030 ERBB2 mutated non-small cell lung cancer cell line and the HCT116 colorectal cancer cell line. We found that the cells containing ERBB2 3’UTR destabilizing constructs show loss of ERBB2 protein (**Fig. 1N, Fig. S1M**), loss of ERBB2 transcript (**Fig. 1P**) and loss of viability (**Fig. 1O**) compared to the wildtype and controls. The morphology of the cells containing the destabilizing constructs desARE3’UTR ERBB2-3 and 30 is different compared to the controls (**Fig. S1L**) in NCI-H2030, while desARE3’UTR ERBB2-1, 3 and 30 show morphological changes compared to controls in HCT116 (**Fig. S1N**). The desARE3’UTR ERBB2 – 3 and 30 degraded ERBB2 transcript in HCT116 cells (**Fig. S1O**).

Taken together, the findings demonstrate that the novel engineered destabilized ARE 3’UTR of ERBB2 effectively degraded ERBB2 transcript and protein expression to kill cancer cells in a clinically relevant model of ERBB2+ overexpressing osimertinib resistant EGFR T790M non-small cell lung cancer, in ERBB2 mutated non-small cell lung cancer, in ERBB2+ expressing colorectal cancer cell line, in difficult to treat ERBB2+ trastuzumab resistant breast cancer cell line, as well as in wildtype ERBB2+ expressing breast cancer.

### Transcriptome analysis show that engineered destabilized ARE 3’UTR ERBB2 degraded ERBB2 and its interactome through the designed nonsense mediated decay pathway

To validate at the transcriptome level that the engineered destabilized 3’UTR ARE of ERBB2 degraded ERBB2 and its interactome through the designed mechanism incorporated in the ERBB2 3’UTR destabilizing constructs, we performed RNA sequencing on the ERBB2 expressing EGFR T790M lung cancer cells NCI-H1975 WT, NCI-H1975 vector and in NCI-H1975 cancer cells containing (desARE3’UTR ERBB2-3 and desARE3’UTR ERBB2-30). We found that desARE3’UTR ERBB2-3 and 30 show strong reduction of ERBB2 and EGFR respectively (**Fig. 2A**). We engineered the constructs to degrade mRNA once the transcript ARE is destabilized through mRNA decay machinery. By gene ontology analysis, we investigated if the mRNA decay mechanism is triggered once the transcript is destabilized and driven by mRNA cap de-adenylating DCP1A promoter. As designed in the constructs, the gene ontology theme of non-sense mediated decay is most significantly enriched GO biological and molecular function with a p value =5.2e-^13^, FDR =5.3e-^10^ and the genes UPF3B and UPF3A which are known regulators of nonsense mediated decay were implicated (**Fig. 2A-C**). Again, ontology theme also identified upregulated 3’UTR mediated translation regulation (p value=1.6e-^11^, FDR= 8.7e-^10^) as part of the mechanism implicated corroborating our engineering design which specifically targeted the ARE on the 3’UTR of ERBB2 (**Fig. 2B**). The GO downregulated theme implicated transmembrane receptor tyrosine kinase (p=0.0008, FDR =0.048) to which ERBB2 which is an EGFR family receptor belongs to (**Fig. 2D**). With UPF3B upregulated (**Fig. 2C**), we searched for premature termination codon on the ERBB2 genes in the destabilized constructs compared to the wild type, we found SNPs that created premature termination codon on exon 4 (UAA) and exon 15 (UGA) of the destabilized constructs (**Fig. 2E**). Indeed, the transcriptome analysis confirmed that these engineered constructs reliably targeted ERBB2 and worked as the engineered constructs were designed to function through mRNA mediated decay driven by destabilizing ERBB2 oncogene ARE driven under de-capping DCP1A promoter to trigger nonsense mediated mRNA decay and de-adenylases to degrade the ERBB2 transcript.

**Figure 2:**
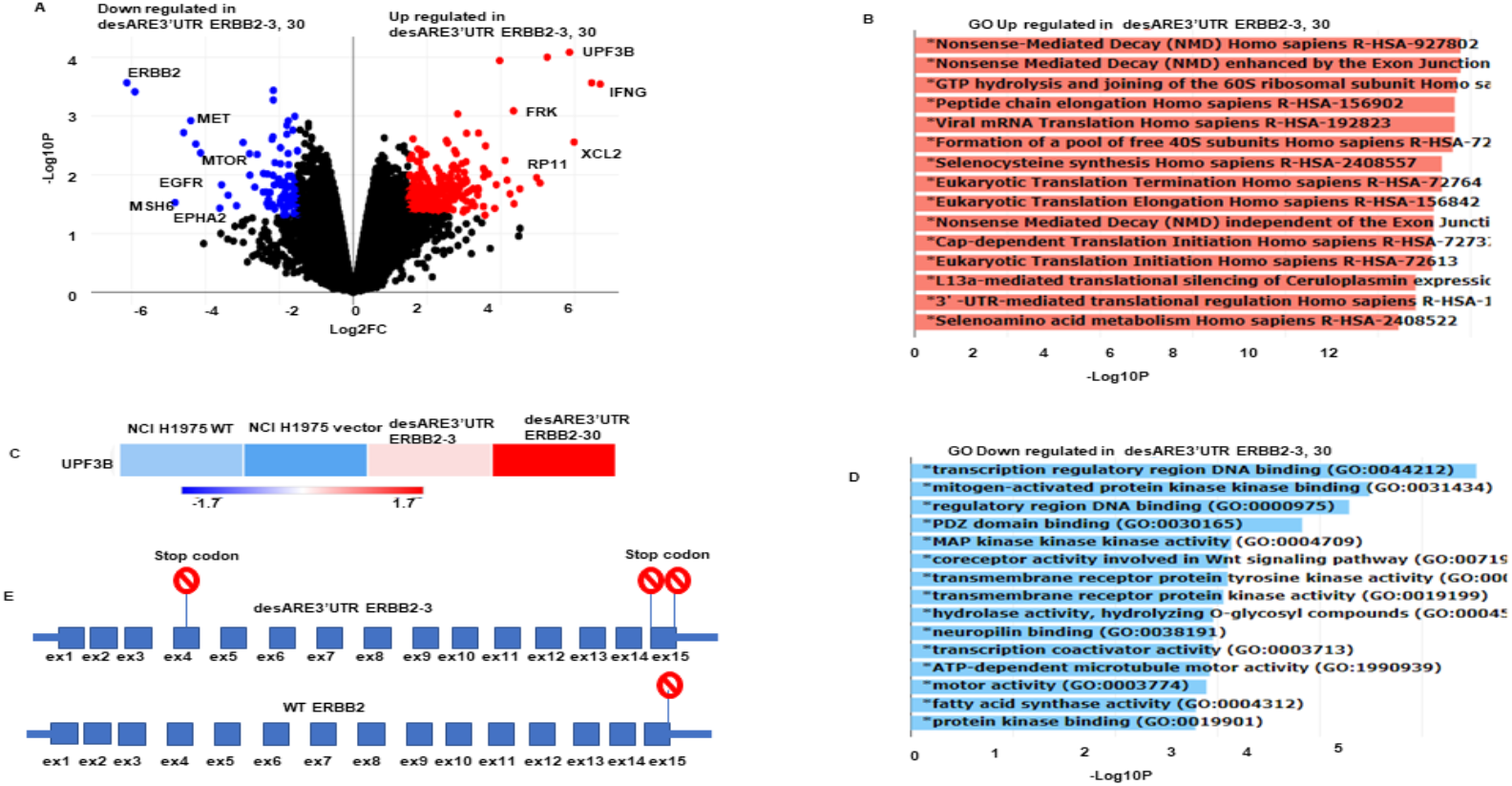
Gene expression analysis of the destabilized 3’UTR ARE of ERBB2 cancer cells show degradation of ERBB2 transcript and worked as designed through upregulated mRNA decay and engineered 3’UTR translational regulation. **A.** The volcano plot shows genes down regulated (blue) and genes upregulated (red) in desARE3’UTR ERBB2-3 and 30 compared to wildtype and vector controls. **B.** The horizontal bar charts show the gene ontology (GO) theme upregulated in desARE3’UTR ERBB2-3 and 30 compared to wildtype and vector controls. **C.** The horizontal bar chart show UPF3B expression in wildtype NCI H1975, vector, desARE3’UTR ERBB2-3 and 30. **D.** The horizontal bar chart shows gene ontology (GO) downregulated in desARE3’UTR ERBB2-3 and 30 compared to wildtype and vector controls. **E.** Schematic depiction of ERBB2 exons comparing stop codon found in the desARE3’UTR ERBB2-3 and 30 compared to wildtype.

### Engineered destabilized 3’UTR ARE loss of ERBB2 is sequence specific and triggers exonuclease XRN1 and CNOT1 to degrade ERBB2 transcript

To ascertain the molecular details of how the degradation of ERBB2 is achieved (**Fig. 1B, 1D, 1F, 1H, IJ, 1L and 1N**). We designed the engineered destabilized 3’UTR ARE of ERBB2 to be driven by the mRNA decapping enzyme DCP1A promoter. To understand if the DCP1A promoter contributed to ERBB2 degradation through mRNA decapping upon the introduction of the engineered destabilizing 3’UTR ARE of ERBB2 into the breast and lung cancer cells, we performed qRT-PCR and showed that DCP1A promoter expression did not change upon destabilization of ERBB2 transcript compared to the controls (**Fig. 3A**). Moreso, DCP1A expression from RNA seq (**Fig. 3B**) show that DCP1A does not increase in the destabilized constructs compared to control. This finding suggests that the engineered destabilized 3’UTR ARE sequence only explains the degradation of ERBB2. To understand the deadenylation proteins involved in this process and if there is a preference of 5’-3’ or 3’-5’ deadenylation. We performed a qRT-PCR on the 5’-3’ deadenylase XRN1 and 3’-5’ deadenylase PARN and CNOT1 both on the BT474 clone 5 and the NCI H1975 containing the destabilized constructs compared to the controls. In the BT474 clone 5 containing the destabilized 3’UTR ARE of ERBB2. We found that the XRN1 expression is significantly elevated in the desARE3’UTRERBB2-3 (**Fig. 3C**) which completely degraded ERBB2 protein in (**Fig. 1F**, **Fig. 1H**). The PARN expression is not elevated (**Fig. 3D**). The CNOT1 expression is significantly elevated in desARE3’UTRERBB2-3 (**Fig. 3E**). We found similar results in NCI H1975 transduced with the destabilizing constructs of the 3’UTR ARE of ERBB2. XRN1 is significantly elevated in desARE3’UTR ERBB2-30 (**Fig. 3F**), also CNOT1 is significantly elevated in the desARE3’UTRERBB2-30 (**Fig. 3H**). The desARE 3’UTR ERBB2-30 showed the most loss of ERBB2 in (**Fig. 1B**). Again, the PARN expression is not elevated in NCI H1975 (**Fig. 3G**) as seen in the BT474 clone 5 (**Fig. 3D**). Finally, to prove the increased CNOT1 protein expression in the destabilized cells, we performed a western blot and show that CNOT1 is highly expressed in the cells carrying the desARE3’UTRERBB2-3,30 (**Fig. 3I-J**). *In-vivo* we found that loss of ERBB2 in animal bearing tumors treated with desARE3’UTR ERBB2-1,3 and 30 and increased expression of UPF3B, XRN1 and CNOT1 (**Fig. S14 A-H**). These results suggest a general mechanism that both 5’-3’ and 3’-5’ deadenylases and nonsense mediated decay are involved in the degradation of the ERBB2 once the 3’UTR ARE of ERBB2 is destabilized.

**Figure 3:**
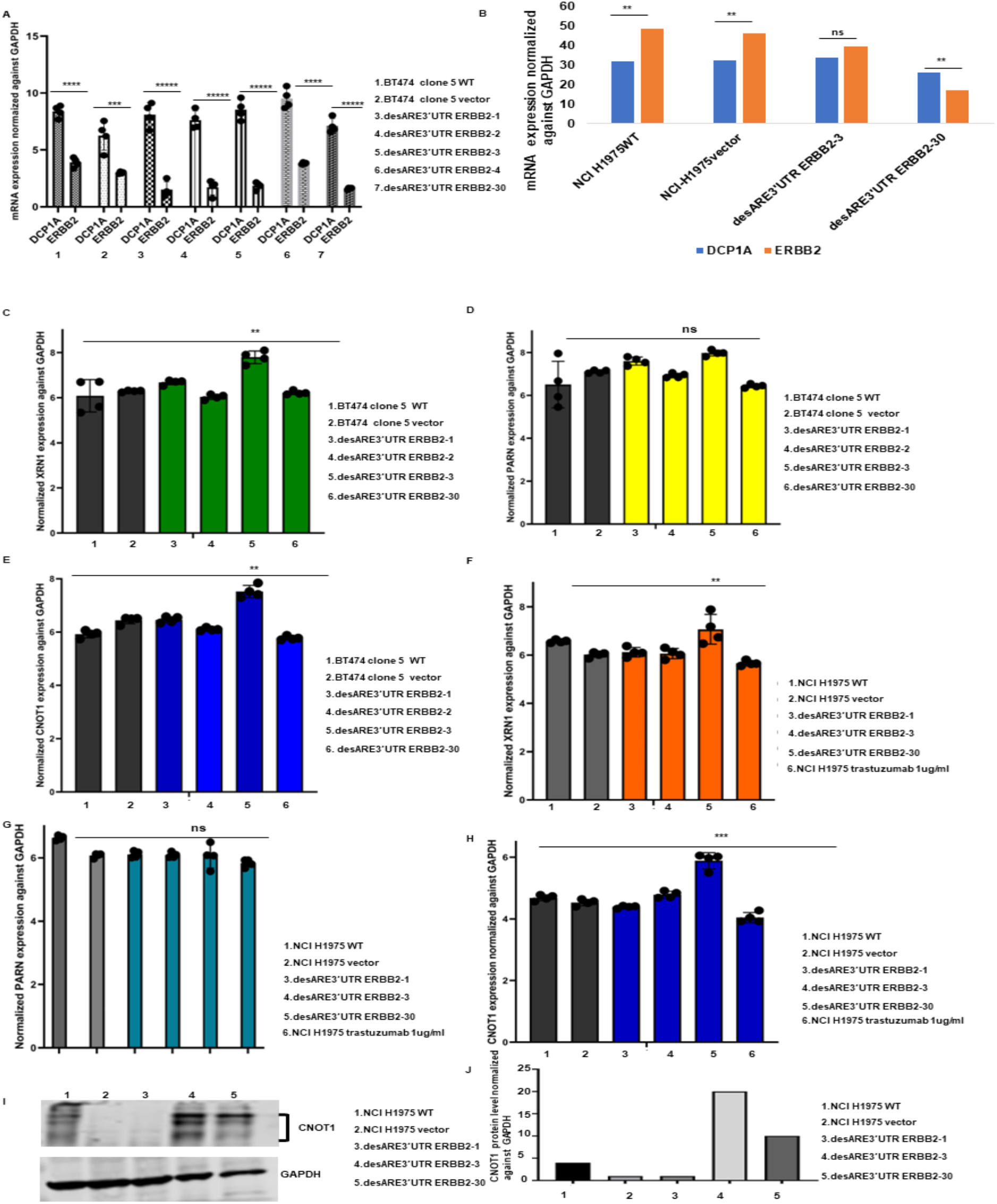
Engineered destabilized 3’UTR ARE of ERBB2 is sequence specific in degrading the ERBB2 transcript through XRN1 and CNOT1. **A.** Bar charts show mRNA expression of DCP1A and ERBB2 normalized against GAPDH in BT474 clone 5 WT, vector, desARE3’UTR ERBB2-1, 2, 3, 4 and 30 (n=2). T-test (****p=0.0001 ERBB2 vs DCP1A BT474 clone 5 WT), (*** p=0.0012 ERBB2 vs DCP1A vector),(*****p=0.00001 ERBB2 vs DCP1A desARE3’UTRERBB2-1), (*****p=0.00003 ERBB2 vs DCP1A desARE3’UTRERBB2-2), (*****p=0.000025 ERBB2 vs DCP1A desARE3’UTRERBB2-3), (****p=0.00013 ERBB2 vs DCP1A desARE3’UTRERBB2-4), (*****p=0.000016 ERBB2 vs DCP1A desARE3’UTRERBB2-30) **B.** Bar charts show the RNA Seq expression of DCP1A in NCI H1975 WT, vector, desARE3’UTR ERBB2-3 and 30. T-test (****p=0.0128 ERBB2 vs DCP1A NCIH1975 WT), (**p=0.011 ERBB2 vs DCP1A vector), (p=ns ERBB2 vs DCP1A desARE3’UTRERBB2-3), (**p=0.001 ERBB2 vs DCP1A desARE3’UTRERBB2-30). **C.** Bar charts show the mRNA expression of XRN1 normalized against GAPDH in BT474 clone 5 WT, vector, desARE3’UTR ERBB2-1, 2, 3, 4 and 30, ** P value =0.002, two tailed t-test (n=2). **D.** Bar charts show the mRNA expression of PARN normalized against GAPDH in BT474 clone 5 WT, vector, desARE3’UTR ERBB2-1, 2, 3, 4 and 30, P value =ns, two tailed t-test (n=2). **E.** Bar charts show the mRNA expression of CNOT1 normalized against GAPDH in BT474 clone 5 WT, vector, desARE3’UTR ERBB2-1, 2, 3, 4 and 30, ** P value =0.001, two tailed t-test (n=2). **F.** Bar charts show the mRNA expression of XRN1 normalized against GAPDH in NCI H1975 WT, vector, desARE3’UTR ERBB2-1, 2, 3, 4, 30 and trastuzumab treated cells ** P value =0.004, two tailed t-test (n=2). **G.** Bar charts show the mRNA expression of PARN normalized against GAPDH in NCI H1975 WT, vector, desARE3’UTR ERBB2-1, 2, 3, 4, 30 and trastuzumab treated cells, P value =ns, two tailed t-test (n=2). **H.** Bar charts show the mRNA expression of CNOT1 normalized against GAPDH in NCI H1975 WT, vector, desARE3’UTR ERBB2-1, 2, 3, 4, 30 and trastuzumab treated cells, P value =0.0002, two tailed t-test (n=2). **I.** Western blot showing CNOT1 and GAPDH protein expression in NCI H1975 WT, vector, desARE3’UTR ERBB2-1, 3, 30. **J.** Bar chart show quantification of CNOT1 protein expression normalized against GAPDH in NCI H1975 WT, vector, desARE3’UTR ERBB2-1, 3, 30.

### Destabilized 3’UTR ARE degradation of ERBB2 in osimertinib and trastuzumab resistant cancer cell lines leads to loss of kinases implicated in breast and lung cancer drug resistance

The signaling cascade of ERBB2 is mediated by kinases and as such the resistance to anti-ERBB2 is concomitant to with the increased activity kinase activities that helps the drug resistant cancers to proliferate and metastasize. Osimertinib EGFR T790M resistance arises due to upregulation of EGFR dependent kinases and EGFR independent pathway. We explored if the destabilization of ERBB2 in this non-small cell lung cancer line does indeed control these implicated kinases and proteins. We found that EGFR, MET, and MAPK kinases implicated in EGFR dependent pathway of osimertinib resistant were all down regulated (**Fig. S2A-B List S2**). More so, the EGFR independent pathways such as MTOR, CNNTB1, EPHA2, NOTCH etc. (**Fig. 2C**) were down regulated upon destabilization of ERBB2. This finding points to the superiority of this approach in controlling drug resistance driven by ERBB2 and its widespread interactome, such that when we control of ERBB2, we controlled all the pathways that are dependent on it.

To ascertain if the destabilized 3’UTR ARE of ERBB2 degradation of ERBB2 led to loss of kinases known to contribute to ERBB2+trastuzumab resistance. We performed a phosphorylation kinase array assay with protein extracts from BT474 clone 5 WT, BT474 clone 5 vector, BT474 clone 5 desARE 3’UTR ERBB2-3 and desARE3’UTR c-MYC 2-3 (a specificity control for ERBB2 destabilization) (**Fig. 1F-H, Fig. 4A-B**). We found that kinases such as AKT1/2/3 pT308, AKT1/2/3 pS473, Chk-2 pT68, c-JUN pS63, P53 pS46,P53pS15,P53 pS392, P70 S6 kinase pT389, P70 S6 kinase pT421/ pS424, PRAS 40 pT246, PYK2 pY402, RSK1/2 pS221/pS227, RSK1/2/3 pS380/pS386,pS377, STAT1pY701, STAT2 pY705, STAT2 pS727,STAT3 pS727, STAT6 pY641 and HSP60 marked with red box in the lower panel of desARE3’UTR ERBB2-3 were all down regulated upon the degradation of ERBB2 by 3’UTR destabilization compared to controls (**Fig. 4A-B**).

**Figure 4:**
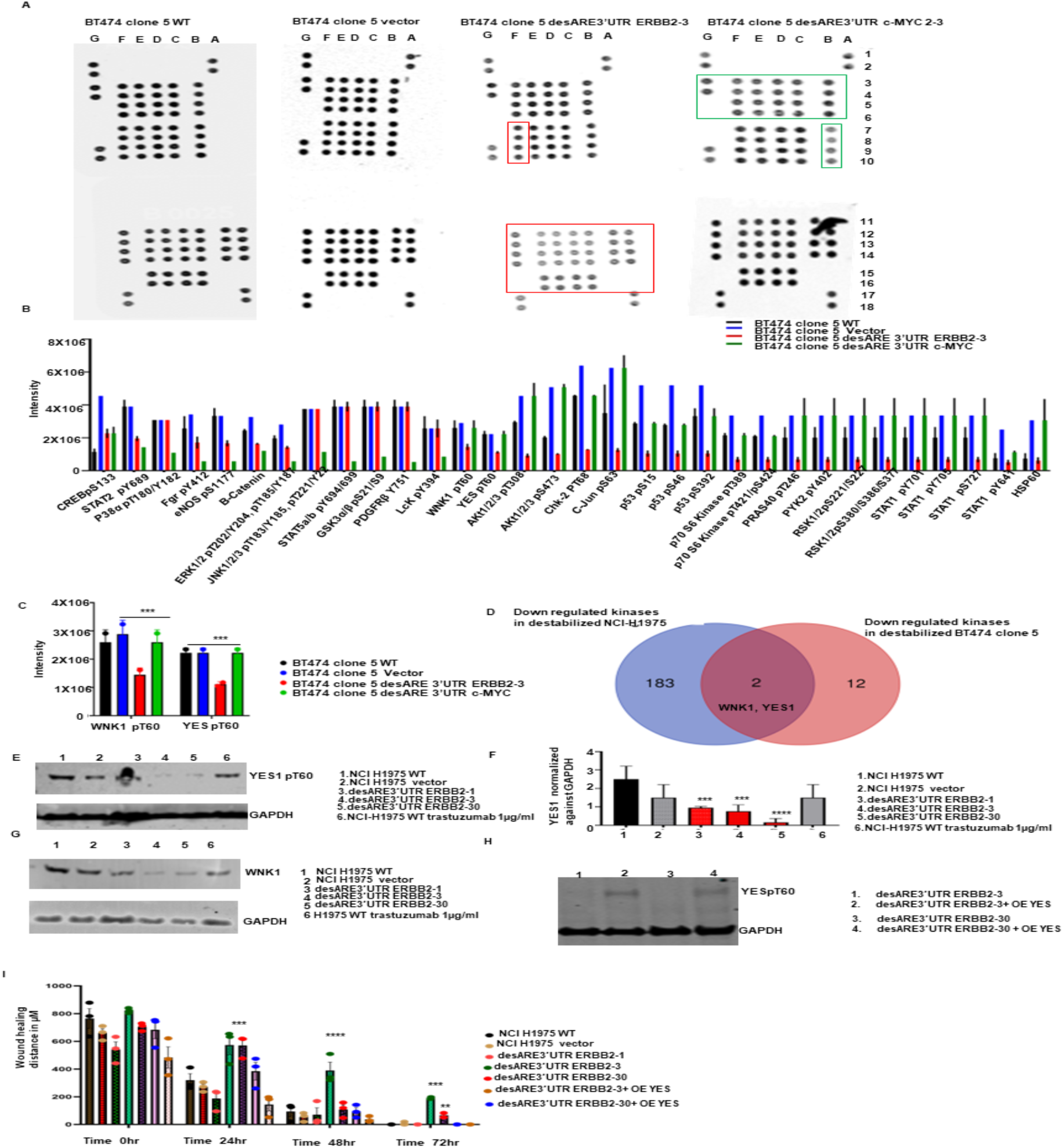
Engineered destabilized 3’UTR ARE of ERBB2 degrades specific kinases implicated in ERBB2+ trastuzumab resistant breast cancer and in ERBB2 expressing Osimertinib resistant EGFR790M non-small cell lung cancer cells. **A.** Dot blots show the phospho kinase array pattern of the BT474 clone 5 wild type, BT474 clone 5 Vector, BT474 clone 5 desARE3’UTR ERBB2-3 and BT474 clone 5 desARE3’UTR c-MYC 2-3. The dot marked in red box show unique kinases down regulated only in BT474 clone 5 desARE3’UTR ERBB2-3 and the dot marked in green boxes show unique kinases downregulated only in BT474 clone 5 desARE3’UTR c-MYC 2-3. Each kinase is spotted twice. **B.** The bar chart shows the intensity of the phospho-kinase array dot blot for the BT474 clone 5 wild type (black bar), BT474 clone 5 Vector (blue bar), BT474 clone 5 desARE3’UTR ERBB2-3 (red bar) and BT474 clone 5 desARE3’UTR c-MYC 2-3 (green bar). **C.** The bar charts show YES1 pT60 and WNK1 pT60 kinases specifically down regulated in BT474 clone 5 desARE3’UTR ERBB2-3 in red bar. Controls BT474 clone 5 wild type (black bar), BT474 clone 5 Vector (blue bar), and BT474 clone 5 desARE3’UTR c-MYC 2-3 (green bar). T-test (*** Pval=0.00413, desARE3’UTRERBB2-3 for YES and WNK1) **D.** Venn diagram show kinases downregulated in destabilized desARE3’UTR ERBB2-3, 30 NCI-H1975 (p=0.02) blue (from RNA Seq), and in destabilized desARE3’UTR ERBB2-3 BT474 clone 5 (red-kinase array) and the shared kinases found downregulated in destabilized ERBB2 in both resistant cell lines WNK1 and YES1 in pink. **E.** Western blot show YES1 and GAPDH expression across the NCI-H1975 wildtype cells, vector, cells containing constructs desARE3’UTR ERBB2-1, 3 and 30 and wild type trastuzumab treated NCI-H1975 cells (n=2). **F.** Bar chart show quantification of the YES1 expression normalized against GAPDH on NCI-H1975 wildtype cells, vector and cells containing constructs desARE3’UTR ERBB2-1, 3 and 30 (n=2). T-test (***Pval=0.0021, desARE3’UTRERBB2-1,3,30) **G.** Western blot show WNK1 and GAPDH expression across the NCI-H1975 wildtype cells, vector, cells containing constructs desARE3’UTR ERBB2-1, 3 and 30 and wild type trastuzumab treated NCI-H1975 cells. **H.** Western blot show YES1 and GAPDH expression across the NCI-H1975 destabilized with constructs desARE3’UTR ERBB2-3 and 30 (lane 1 and 3) and over expressed with YES1 in lane 2 and 4. **I.** Bar chart show the wound healing closing of NCI H1975 wildtype, vector, desARE3’UTRERBB2-1, 3,30 and desARE3’UTR ERBB2-30,30 overexpressed with YES1 from 0hr to 72hrs. At 24hrs *** p=0.00025 desARE3’UTRERBB2-3,30. At 48hrs ****p=0.000018 desARE3’UTR ERBB2-3 and at 72hrs *** p=0.0003 desARE3’UTRERBB2-3 and **p=0.002 desARE3’UTRERBB2-30.

These kinases downregulated has been implicated in mediation of trastuzumab resistance in breast cancer, both from clinical trials and in cancer cell lines (***28-31***). Secondly, the result revealed ERBB2 is a master regulator controlling vast array of kinases and transcription factor and that our novel developed technology can control not only the ERBB2 but as well all the signaling cascade and interactome associated with it. We also explored kinases that were lost in desARE3’UTR ERBB2-3 samples but whose expression is relatively similar both in wildtype, vector control and desARE 3’UTR c-MYC 2-3; a specificity control for ERBB2 destabilization, we found the kinases YES pT60 and WNK1 pT60 (**Fig. 4A-B**, **Fig. 4C**) marked with red box in upper panel of desARE3’UTR ERBB2-3. To find the universal common kinases that are mediate drug resistance both in osimertinib resistant EGFR T790M lung cancer and trastuzumab resistant breast cancer that are downregulated upon our destabilization of ERBB2. We triaged all the kinases downregulated in ERBB2 destabilized NCI-H1975 (**List S2**) found by RNA Seq with kinases downregulated in ERBB2 destabilized BT474 clone 5 (**Fig. 4A-C**) found by kinase array. Indeed, we found that WNK1 and YES1 is the common shared kinase by these two cancer cell lines that mediates drug resistance in them amongst other (**Fig. 4D**).

To assess the specificity of the kinases downregulated through the destabilization of the ARE 3’UTR of ERBB2 is unique to ERBB2 and not to other oncogenes such as c-MYC. We performed phosphorylation kinase array assay on protein extract of the BT474 clone 5 destabilized with desARE3’UTR MYC 2-3. We found in (**Fig. 4A-B**), the last panel marked with green box, that the kinases lost by the destabilization of c-MYC is unique to c-MYC and that kinases lost by destabilization of ERBB2 is unique to ERBB2, proving surprisingly that the destabilization of the ARE 3’UTR of oncogenes is unique and specific and a targeted molecular therapy.

To validate the loss of YES1 pT60 and WNK1 upon the destabilization of the ARE of 3’UTR of ERBB2 (**Fig. 4A-D**). We performed western blot on desARE3’UTR ERBB2 samples compared to the controls. We found that YES1 pT60 protein expression were decreased in the desARE3’UTRERBB2-3 and 30 compared to the controls across multiple ERBB2 driven cancer types such as NC-H1975 (**Fig. 4D, 4E, 4F**). The engineered destabilized desARE3’UTRERBB2-1,3,30 degraded WNK1 (**Fig.4G**). Taken together, we demonstrate that by destabilizing ERBB2, that we controlled and downregulated YES1 and WNK1 that are well known causes of drug resistance in ERBB2+ trastuzumab drug resistant and in osimertinib resistant EGFR T790M lung cancer (***28-31***).

In a gain of function experiment, we overexpressed YES1 using a construct developed by Rosenbluh J et al in (***32***), in NCI H1975 desARE3’UTR ERBB2-3 and 30 cells that we degraded and reduced ERBB2 and YES1. We confirmed the overexpression of YES1 by western blot (**Fig. 4H**). Next, we performed wound healing assay comparing the wildtype control, vector, the desARE3’UTR ERBB2-1,3,30 and the desARE3’UTRERBB2-3, 30 with overexpressed YES. We found that the (**Fig. 4I, Fig. S11**) desARE3’UTRERBB2-3, 30 OE YES closed their wound at similar time point as the controls. The cells bearing the destabilized desARE3’UTRERBB2-3,30 was unable to close their wounds due to lack of ERBB2 and YES (**Fig. 4I, Fig. S11**). This finding confirms ERBB2 and it’s downstream YES kinase control resistance in NCI H1975 and that this novel approach we have developed is superior in controlling ERBB2 pervasive oncogene signal and their downstream YES kinases that are involved in drug resistance which osimertinib and trastuzumab can’t control.

### The engineered destabilized ARE 3’UTR of ERBB2 outperforms ShRNAi and genome scale CRISPR in controlling ERBB2 transcript in ERBB2 expressing colorectal cancer HCT116

To evaluate the performance of the engineered destabilized ARE 3’UTR of ERBB2 in controlling ERBB2 transcript and downstream kinases in comparison to ShRNAi and genome scale CRISPR. We analyzed the public available database for ShRNAi and CRISPR (***33-35***) in HCT116, a colorectal carcinoma cell, a ERBB2 expressing cancer. We found (**Table 1**) that the ShRNAi and CRISPR across multiple studies were unable to downregulate ERBB2 transcript in HCT116 and its associated kinases, whereas the desARE3’UTR ERBB2 degraded ERBB2 transcript in this study (**Fig. S1 N, O**) as well as YES1 (**Fig. 4I**). Hart T et al Cell 2015 (***34***) suggested that lack of TP53^S241F^ EGFR dependency in HCT116 prevented erlotinib and the ERBB2 and EGFR sgRNA from controlling ERBB2 in this cell line HCT116 KRAS ^G13D^. The engineered destabilized ARE 3’UTR of ERBB2 is agnostic to these mutations and reliably controlled and degraded ERBB2 transcript in this cancer. This finding strongly suggests that the destabilization of stable oncogene 3’UTR ARE is superior to RNAi and CRISPR in certain genetic mutation settings.

**Table 1.** Comparison of ShRNAI, CRISPR and mRNA 3’UTR destabilized in targeting HER2 in colorectal carcinoma HCT116.

| **Transcript perturbation method** | **Publication** | **Target** | **Cell line** | **Outcome** | **Associated kinases outcome** |
| --- | --- | --- | --- | --- | --- |
| Genomewide ShRNA | Vizeacoumar F et al Mol Syst Biol 2013 | EGFR, ERBB2 | HCT116 | ERBB2 not downregulated | YES1 not downregulated |
| Genome wide CRISPR KO | Hart T et al Cell 2015 | EGFR , ERBB2 | HCT116 | EGFR and ERBB2 not downregulated |  |
| Genome wide CRISPR KO plus shRNA | Martin T et al, Cell Rep 2017 | EGFR , ERBB2 | HCT116 | EGFR and ERBB2 not downregulated | YES1 not downregulated |
| desARE 3'UTR ERBB2-1,3 | Awah CU et al 2022, This Publication | EGFR , ERBB2 | HCT116 | ERBB2 downregulated | YES1 downregulated |

### Engineered destabilized ARE3’UTR of ERBB2 reprogrammed lung cancer ERBB2 gene expression towards the normal lung epithelial gene expression pattern

To understand if the destabilization of the ERBB2 transcript by the 3’UTR engineering, restored the lung genetic program towards normal lung epithelial cells. We compared the ERBB2 gene expression profile of wildtype NCI H1975, vector, desARE3’UTR ERBB2-3,30 and BEAS-2B, a normal human lung epithelial cell line (E-MTAB-4729, **List. S3**). We found that the degrading of the ERBB2 transcript by destabilized ARE 3’UTR changed the cancer ERBB2 gene expression almost to near normal lung epithelial cells gene expression program (**Fig. 5A, List. S3**). We extended this data to compare if the destabilization of the ERBB2 transcript and protein in cancers does reprogram the gene expression of cancers towards normal human lung gene expression pattern. To do this, we obtained the RNA Seq of diverse normal human lung individuals (normal Chinese female non-smoker, SRR1797221, normal Caucasian male, normal female (3) and normal male (3) ENCODE data **List. S3**). Indeed, we found in (**Fig.5B**) that the desARE3’UTR ERBB2-30 reprogrammed the cancers ERBB2 gene expression to normal pattern as in healthy humans. This data strongly suggests that 3’UTR destabilization of oncogene transcript is safe and achieves the restoration of normal gene expression patterns.

**Figure 5:**
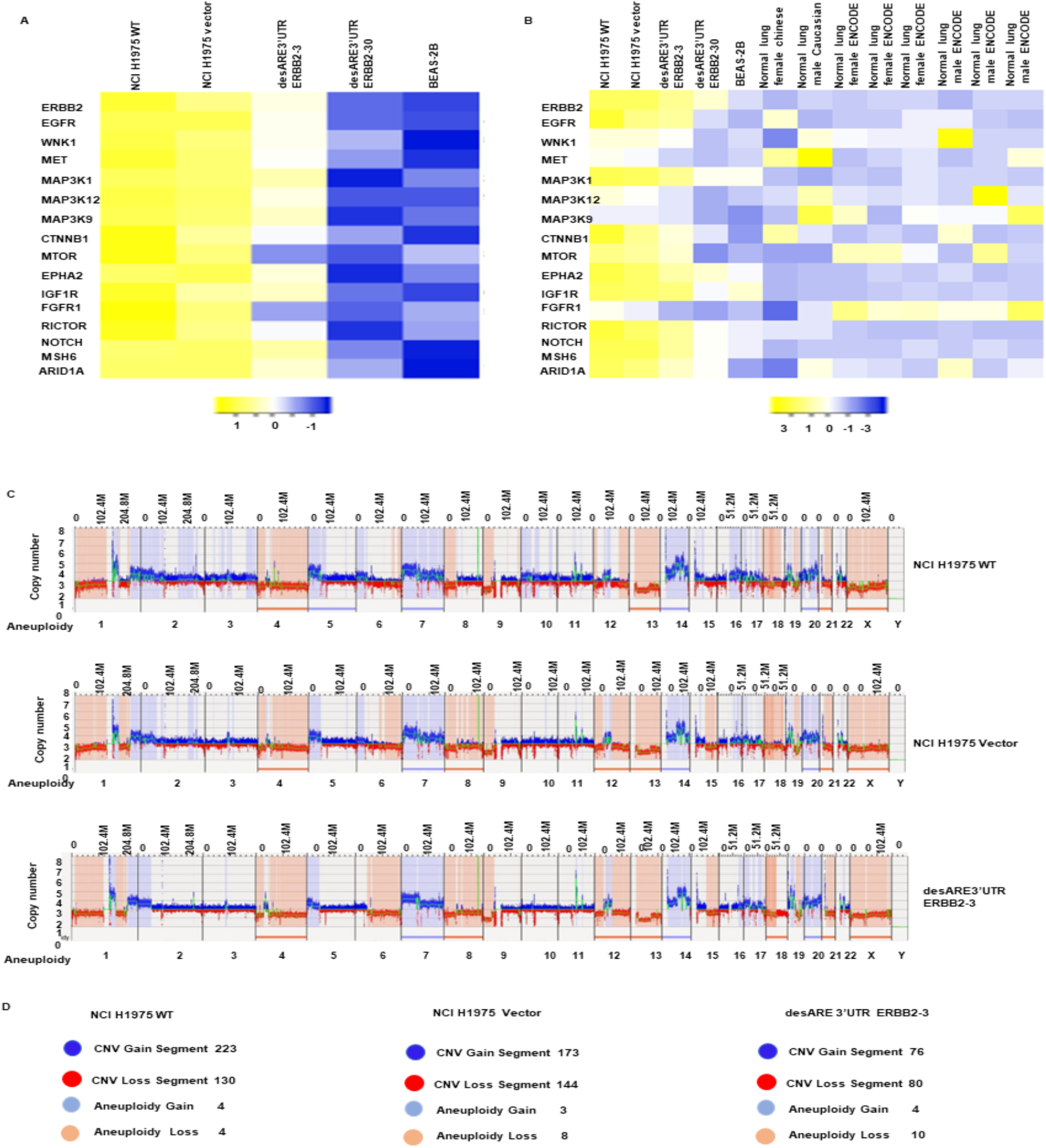
Engineered destabilized 3’UTR ARE of ERBB2 degradation of ERBB2 transcript reprogramed lung cancer ERBB2 gene expression toward normal lung gene expression profile. **A.** Heat map shows the ERBB2 gene expression profile in wildtype NCI-H1975, vector, desARE3’UTR ERBB2 -3, 30 compared to the BEAS-2B normal human lung epithelial cells. **B.** Heat map shows the ERBB2 gene expression profile in wildtype NCI-H1975, vector, desARE3’UTR ERBB2 -3, 30 compared to the BEAS-2B normal human lung epithelial cells line and normal lungs from a Chinese woman non-smoker, normal lung from a Caucasian male, 3 normal female lungs from ENCODE and 3 normal male lungs from ENCODE. **C.** Genome browser view of whole genome de novo variant assembly between NCI-H1975 wildtype, vector and desARE3’UTR ERBB2-3. **D.** Schematic representation of the genomic alteration and their quantification in wildtype NCI-H1975, vector, desARE3’UTR ERBB2 -3. Dark Blue color (CNV segment gain), red (CNV segment loss), Light blue (Aneuploidy gain), Light Orange (Aneuploidy loss) (n=2).

### Genome wide optical genome mapping show that engineered destabilized ARE 3’UTR ERBB2 de novo and rare variants assembly are similar with parental cell line

To rule out that our engineered destabilization of the 3’UTR ARE of ERBB2 did not cause severe genome alterations and genome rearrangements in the engineered cells compared to the wildtype and control cells. We used the ultrasensitive technique of optical genome mapping (***36***) and found that the cells carrying the constructs desARE3’ERBB2-3 showed almost the same de novo and rare variant assembly compared to the wild type cells and the vector (**Fig. 5C, Fig. S9A, 10A**). We found that the desARE3’UTR ERBB2-3 has lesser copy number variation segment gain, lesser copy number variation segment loss and more aneuploidy loss compared to the wildtype and vector. This result can be interpreted as the destabilizing constructs desARE3’UTR ERBB2 is safe for the cells and had lesser deleterious changes than the wildtype and vector control (**Fig. 5D**). Aneuploidy is a major cause of chromosomal instability and cancer. It is surprising to find in this results that the engineered constructs destabilized ERBB2 and led to a greater loss of aneuploidy in the cancers carrying the constructs compared to the controls. These results suggest that destabilized 3’UTR ARE degradation of ERBB2 has anti-cancer activity amongst which is the reduction of cancer cells aneuploidy.

### Engineered destabilized ARE 3’UTR of ERBB2 significantly decreased tumor volume *in vivo* with no toxicity or adverse effects

To validate our findings that the engineered destabilized 3’UTR ARE of ERBB2 degrades ERBB2 in-vivo and reduces ERBB2 driven tumor growth and volume in a mouse tumor bearing model. We implanted 25 female mice with 5million NCI H1975 cells on the flank and after 35 days, huge tumors engrafted (**Fig. 6A**). On the day 36 (**Fig. 6A**), we randomized the mice into 5 different group namely: wildtype, vector, desARE3’UTRERBB2-1, desARE3’UTRERBB2-3 and desARE3’UTRERBB2-30 with the groups having equal representation of tumor size. We then administered the vector, the constructs desARE3’UTRERBB2-1, 3 and 30 at 20µg per dose given intraperitoneal 12hrly for 9 days (**Fig. 6A**). Upon each administration, the mice were weighed, and tumor size measured with calipers both length and width recorded. And the body condition score of the mice were recorded too. We plotted the tumor volumes comparing the wild type and the mice that received the vector, desARE3’UTRERBB2-1, 3 and 30. We found that (**Fig. 6B, C**) the tumor volumes were significantly reduced in the mice bearing tumors that received the engineered constructs desARE 3’UTR ERBB2-1, 3 and 30 compared to the wildtype and vector respectively and that there were no significant differences between tumor volumes in the wildtype and the vector (**Fig. 6B, C**). This finding strongly validates in-vivo the findings already established here in that the engineered destabilized 3’UTR ARE of ERBB2 degraded ERBB2 and its interactome and impaired cancer cell growth, size, and migration.

**Figure 6:**
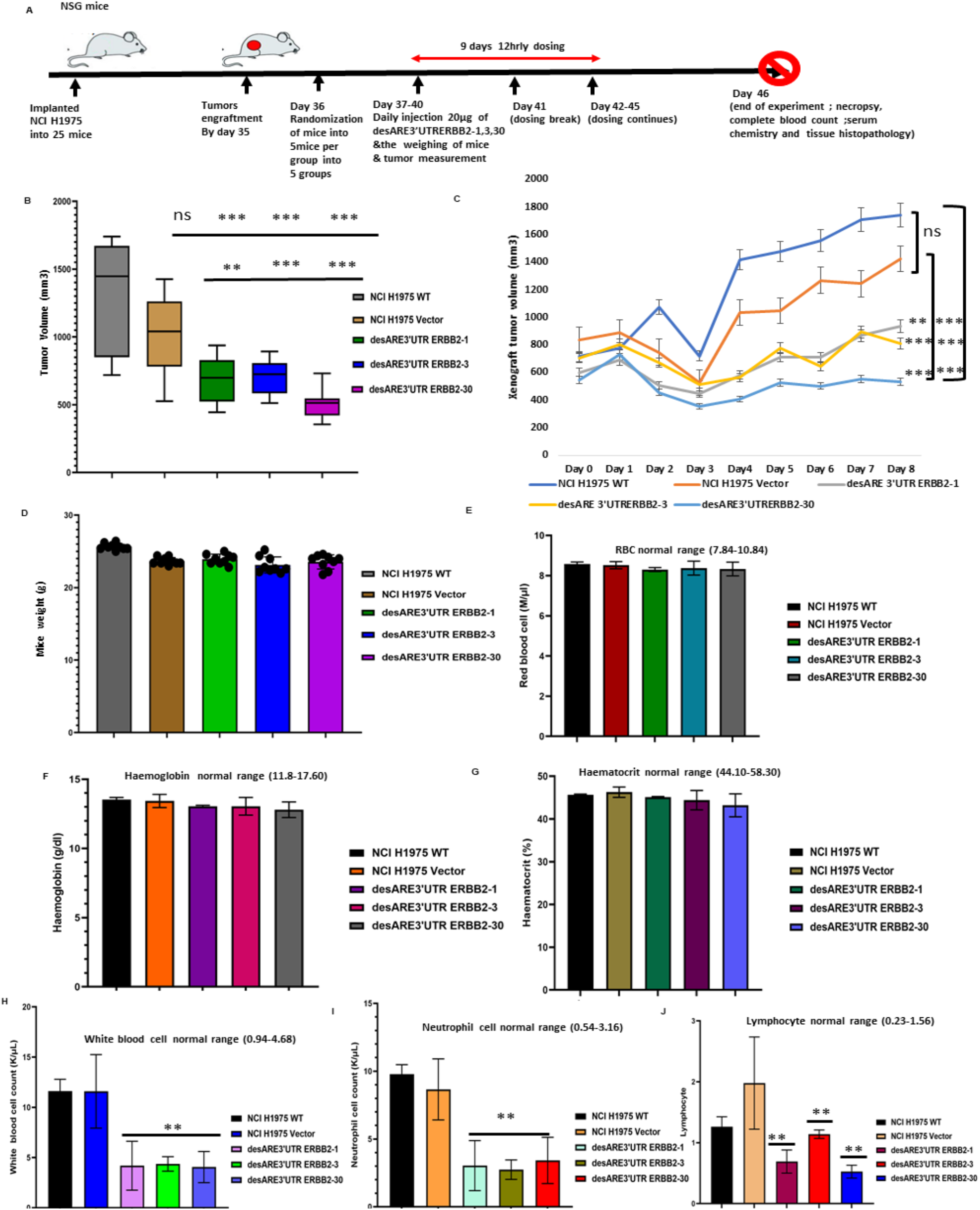
Engineered destabilized ARE 3’UTR of ERBB2 significantly decreased tumor volume in-vivo with no toxicity or adverse effects. **A.** Schema depicts outline of animal experiment.25 NSG female mice were implanted with 5million NCI H1975 as flank xenograft, after 35 days post implantation, huge tumor engrafts. Day 36 post implantation, we randomized the tumors for equal size distribution. Treatment groups were administered 20µg IP 12hrly of engineered constructs for 9days with daily weight and tumor size measurement. Day 46 post implantation, control groups tumor sizes have grown so enormous that mice were euthanized full blood count, electrolyte, necropsy, and histopathological analysis were performed by expert pathologist. **B.** Box plot show tumor volumes in NCI H1975 WT (gray), vector (light brown), desARE3’UTRERBB2-1(green), desARE3’UTRERBB2-3 (blue), desARE3’UTRERBB2-30 (purple). Two tailed paired test, P-value ns= WT vs vector, WT vs desARE3’UTRERBB2-1,3, 30 (***0.0026, ***0.0077, ***0.004), Vector vs desARE3’UTRERBB2-1,3,30 (**0.0251, ***0.0088, ***0.0031). **C.** Line plot show daily tumor growth comparing NCI H1975 WT (dark blue), Vector (brown) and desARE3’UTRERBB2-1 (gray), desARE3’UTRERBB2-3 (yellow), desARE3’UTRERBB2-30. Two tailed paired test, P-value ns= WT vs vector, WT vs desARE3’UTRERBB2-1,3, 30 (***0.0026, ***0.0077, ***0.004), Vector vs desARE3’UTRERBB2-1,3,30 (**0.0251, ***0.0088, ***0.0031). **D.** Bar charts show weight of mice bearing tumor NCI H1975 (WT-gray), vector (brown), and mice bearing tumors treated with desARE3’UTRERBB2-1(green), desARE3’UTRERBB2-3 (blue), desARE3’UTRERBB2-30 (purple). **E.** Bar charts shows red blood cell count of mice bearing tumor NCI H1975 WT (black), vector (red), and mice bearing tumors treated with desARE3’UTR ERBB2-1 (green), desARE3’UTRERBB2-3 (blue), desARE3’UTR ERBB2-30 (gray) **F.** Bar charts show hemoglobin count of mice bearing tumor NCI H1975 WT (black), orange (vector), and mice bearing tumors treated with desARE3’UTR ERBB2-1 (pink), desARE3’UTR ERBB2-3 (magenta), desARE3’UTR ERBB2-30 (gray). **G.** Bar charts show hematocrit level of mice bearing tumor NCI H1975 WT (black), vector (beige), and mice bearing tumors treated with desARE3’UTRERBB2-1 (green), desARE3’UTRERBB2-3 (purple) and desARE3’UTRERBB2-30 (light blue). **H.** Bar charts show white blood cell count of mice bearing tumor NCI H1975 WT (black), vector (blue) and mice bearing tumors treated with desARE3’UTRERBB2-1(pink), desARE3’UTR ERBB2-3 (green), desARE3’UTR ERBB2-30 (blue) **p=0.0065. **I.** Bar charts show neutrophil cell count of mice bearing tumor NCI H1975 WT (black), vector (yellow), and mice treated with desARE3’UTRERBB2-1(light green), desARE3’UTRERBB2-3 (green) and desARE3’UTRERBB2-30 (red). ***p =0.0032 **J.** Bar charts show lymphocyte cell count of mice bearing tumor NCI H1975 WT (black), vector (light orange), and mice treated with desARE3’UTRERBB2-1(magenta), desARE3’UTRERBB2-3 (red) and desARE3’UTRERBB2-30 (blue). **p =0.0002 desARE3’UTRERBB2-1,30. **p=0.0021 desARE3’UTRERBB2-3.

Next, we analyzed the weight of the mice as an early pointer of toxicity. We found that there is no difference between the wildtype, vector and the mice receiving the engineered desARE3’UTRERBB2-1, 3 and 30 (**Fig. 6D**). This finding shows that while the tumor volumes in the controls were increasing the tumor volume, the mice that received the engineered constructs had their tumor volume reduced and they gained weight by feeding well which points to absence of toxicity.

To prove at the molecular level that our novel technology had no adverse effects or toxicity on the mice that received them. We performed a complete blood count and complete blood chemistry analysis on the controls (wildtype and vector tumors) and the desARE3’UTRERBB2-1,3 and 30 mice (**List. S4, S5**). We found that there is no change on the red blood cell count (**Fig. 6E**), hemoglobin (**Fig. 6F**) and hematocrit (**Fig. 6G**), this point to the fact that the engineered constructs are safe and does not cause red blood cell dyscrasias. We analyzed the white blood cells, neutrophils, and lymphocyte. We found that white blood cells, neutrophils and (**Fig. 6H-J**) were significantly elevated in the wild type and vector mice, where as the desARE3’UTRERBB2-1,3 and 30 restored the white blood cells, neutrophils, and lymphocytes to the normal levels. This finding confirms that the engineered constructs while reducing the tumor volume by degrading ERBB2 restores the white blood cells, neutrophils, and lymphocyte to the normal level. The high level of the white blood cells, neutrophils and lymphocyte in the controls shows that the immune cells in those mice are elevated due to the tumor size increase.

Next, we studied the complete blood chemistry of the tumor bearing control mice and those receiving the desARE3’UTRERBB2-1,3 and 30. In the kidney function analysis, we found no increase in blood urea nitrogen and creatinine levels (**Fig. S12A-B**). The liver function test shows no increase in albumin (**Fig. S12E**) and globulin (**Fig. S12F**) in controls and tumors bearing constructs. The alkaline phosphatase levels are low across the control and tumor bearing mice treated with the engineered destabilized constructs (**Fig. S12C**). The aspartate amino transferase level (**Fig. S12D**) was elevated across all mice bearing tumors except for the mice receiving the desARE3’UTR ERBB2-1. The blood glucose (**Fig. S12H**), cholesterol (**Fig. S12I**), triglycerides (**Fig. S12J**) and total bilirubin (**Fig. S13D**) levels are normal between the controls and the tumor bearing mice receiving the engineered constructs. These results strongly shows that there is no abnormal liver, bone, pancreas, and gall bladder function in the mice receiving the engineered destabilized constructs.

Furthermore, we examined the blood electrolyte levels comparing the controls and the tumor bearing mice receiving the constructs. We found that the calcium (**Fig. S12G**), blood CO2 (**Fig. S12K**), sodium (**Fig. S13A**), potassium (**Fig. S13B**) and chloride (**Fig. S13C**) levels were all normal.

Taken together, we have validated in-vivo that the engineered destabilized 3’UTR ARE of ERBB2 significantly reduced tumor volume in a deadly drug resistant tumor model with no abnormal blood, liver, kidney, bone, pancreas, gall bladder and electrolyte imbalance. These findings strongly suggest that the therapy is safe and can be rapidly translated to human cancers. Finally, we stained the xenografts slides of WT mice tumors, vector and tumors treated with desARE3’UTRERBB2-1, 3 and 30 with antibodies against ERBB2, CNOT1, UPF3B and XRN1 (**Fig. S14 A-H**). The results shows that ERBB2 protein expression is significantly lost in mice that received the constructs compared to the controls (**Fig. S14A-B**). CNOT1 expression is significantly upregulated in desARE3’UTR ERBB2-3 **(Fig. S14C -D**) and UPF3B and XRN1 are significantly upregulated in desARE3’UTRERBB2-1,3 and 30 compared to the controls (**Fig. S14 E-F, G-H**). This finding strongly validates our in-vitro results across multiple cancers and validated the biology of design and the mechanistic of the engineered constructs.

## Discussion

To our knowledge, this is the first report of a demonstration of effective targeting of an oncogene by introducing destabilizing motifs into its 3’UTR resulting in destabilization of its transcript and degradation of its protein, and consequent death of the cancer cells in *vitro* and inhibition of *in vivo* tumor growth. We have developed a novel technology by engineering the destabilized 3’UTR ARE that reliably degrades oncogenic transcript and protein expression and their downstream kinases that cause cancer drug resistance. The findings provided in this study show that the deadly cancer drug resistance driven by pervasive oncogenic signals can indeed be controlled by this technology. This new technology demonstrates the ability to modulate levels of oncogenic transcripts by replacing the stabilizing with the destabilizing elements in the 3ʹ UTR in difficult to treat cancers. This work represents a technical advancement and a new strategy for reducing oncogene expression and their interactome. We provide proof-of-principle evidence in this study with findings from diverse cancer models and established a new paradigm which has opened the doors to applying this technique to control any transcript of interest in temporal and spatial scale.

This technology is versatile and generally applicable to controlling diverse oncogenes and cellular transcript across any tissue of interest. We show in this study, that we can control ERBB2 across wide range of cancers namely NCI-H1975 ERBB2 expressing osimertinib resistant EGFR T790M non-small cell lung cancer, wild type ERBB2+ breast cancer BT474, BT474 clone 5 ERBB2+trastuzumab resistant breast cancer, NCI-H2030 ERBB2 mutated non-small cell lung cancer and ERBB2 expressing HCT116 colorectal cancer. The technology is tissue agnostic, but oncogene targeted.

The breakthrough in the functionality of this novel method is the accuracy of biology by design based on universal gene regulatory principles. We designed the constructs to have the destabilized sequences of the AU rich element as described in the methods in which all the possible stabilizing elements were changed (***23-24***) and that this destabilized AU rich element will be driven by mRNA decapping protein DCP1A which will specifically upregulate the mRNA decay pathway and trigger the de-adenylases CNOT1 and XRN1 to degrade the transcript of interest. Also integrated in the design is the sustained slow decay rate in the constructs. Some nucleotides were modified as described in (***26***) to achieve a slow decay rate destabilizing construct. Our experimental results show that the optimal constructs worked as designed in destabilizing the ERBB2 transcript, degrading the transcript through mRNA de-adenlylases CNOT1 and XRN1 and by nonsense mediated decay. The destabilizing constructs integrated into the genome and outcompeted the endogenous ERBB2 and degraded its transcript and protein.

The destabilization of oncogene transcript we have described here is versatile. We have shown that the approach works in the oncogene of interest regardless of the tissue. We demonstrated that ERBB2 can be targeted and controlled with tumor death across many tissues and in diverse biological background namely in EGFR T790M lung cancer, in ERBB2+ wild type breast cancer, in ERBB2+ trastuzumab resistant breast as well as in ERBB2 mutated lung cancer and ERBB2 expressing colorectal cancer. Our results suggest that this technology has the potential to be superior in controlling cancer chemoresistance driven by oncogene and its kinases. In osimertinib and trastuzumab resistant cancers, we show that once we control the oncogenes, we controlled the kinase signaling pathways and interactome through which they function to become resistant. For example, in ERBB2 overexpressing EGFR T790M lung cancer, we controlled and degraded ERBB2 and as well degraded EGFR, MET and YES1 kinases that are known to drive this tumor resistance to drug. Same finding was made in ERBB2+ trastuzumab resistant breast cancer, in that the control of ERBB2 lead to loss of kinases such as WNK1 and YES1 implicated in this cancer mechanism of resistance.

The optimal constructs show difference in the rate of degradation of ERBB2. This is explained by the method of cloning of the constructs and the evolutionary pressure introduced in the destabilizing constructs by the E. coli in which they were cloned into. The desARE3’UTR ERBB2-1, 2, 3 and 4 were obtained by cloning the constructs into recombinant efficient E. coli (**Fig. S7**). Of these 4 constructs, desARE3’UTR ERBB2-3 and desARE3’UTR ERBB2-1 degraded both transcript and protein in most ERBB2 cellular settings both in wildtype and drug resistant settings. The desARE3’UTR ERBB2-2 down regulated transcript and protein in the native ERBB2 WT setting but did not perform optimally in the ERBB2 drug resistant setting. The desARE3’UTR ERBB2-4 was the least performing. We extended our work by using Gibson assembly to clone the constructs in a recombinant deficient E. coli. We obtained the desARE3’UTRERBB2-30 (**Fig. S8**). The desARE3’UTRERBB2-30 is effective as desARE3’UTRERBB2-1 and 3 in degrading ERBB2 across multiple ERBB2 cancer types. The desARE3’UTR ERBB2-30 has little to no mutation in the engineered destabilized ARE that we designed marked in red underline bars in (**Fig. S4)**. The deARE3’UTR ERBB2-1 and 3 contains some point mutation introduced by the recombinant efficient E. coli. Pointing to the fact that bacterial evolutionary pressure further selected and enhanced our construct’s function. It is worthwhile to mention that we designed and engineered the destabilizing constructs based on the BT474 wild type ERBB2 3’UTR ARE. Yet, they work across various ERBB2 expressing cancers from diverse biological organs. The constructs showed anti-cancer activity such as impaired cancer cells viability, reduced migration and induction of apoptosis and increased caspase 3/7, cleaved caspase 3 and caspase 9 expression.

We found that our constructs do not kill normal breast epithelial cells MCF10A (**Fig. S2H**). This proof of principle findings strongly suggest that our destabilizing constructs does not affect normal cell but cancer specific and on target, which is very promising and only works in cancers carrying the stabilizing ERBB2 elements. We show that destabilizing ERBB2 does not affect c-MYC and its kinases (**Fig. 4A**). Our results and finding are now validated in an animal tumor bearing model, where tumor volume is significantly reduced by the engineered destabilized constructs, and no abnormal vital organ functions or electrolyte imbalance were found (**Fig. 6, Fig. S12, S13**). Technologies such as CRISPR (***37***), RNAi (***38***), the degron system (***39***) and PROTAC (***40***) are competitors to the findings presented in this study. CRISPR has been used to knockout gene expression with high efficiency of up to 90% loss of protein. However, CRISPR has remarkably high off target effects and can cause unwanted chromosomal rearrangements as has been shown. Regarding RNAi, even though it can interfere and silence the expression of a protein, it has also off target effects in order of magnitude greater than CRISPR. The degron system has been effective in mediating degradation of proteins, however, the system relies on inducibility by auxin or similar molecules to initiate it functions. The degron degradation is leaky and requires high levels of auxin (***41***). Lastly, for PROTAC which targets proteins that has already been made, the limitation of PROTAC is highly toxic, costly to make, targets only intracellular proteins, cannot target membrane protein. This engineered destabilized 3’UTR ARE mediated degradation of transcript offers a unique and novel approach to target oncogenic transcript at their untranslated region which has been difficult to study. The engineered destabilized 3’UTR ARE are easy to synthesize as gblocks, cheap and easy to clone and they are very specific to target oncogenes in cancer cells but do not degrade proteins in normal cells and the therapeutic degradation of transcript can be achieved at a very low dose of nanogram of vectors containing the destabilizing constructs. This makes the technology very attractive. We have shown that in certain genetic settings of cancer as in ERBB2 HCT116, that we are superior to CRISPR, RNAi and shRNA in a head-to-head comparison. This is because these technologies show genetic dependencies whereas our technology is agnostic to these dependencies and degrades transcript once the stabilizing elements are destabilized (**Table 1**). We show that our constructs reprogram the ERBB2 genetic programs toward the normal individual ERBB2 gene program from diverse ethnicity (**Fig. 5A-B**). By optical genome mapping, the de novo, rare variants and genomic alterations is not different between the wild type, vector and the ERBB2 mRNA destabilized cancer (**Fig. 5C-D**). This evidence strongly suggest that our constructs are safe for the cells. In-fact, our data in (**Fig. 5D**) show that the desARE3’UTRERBB2-3 also lead to loss of aneuploidy in cancer cells, aneuploidy is a major cause of chromosomal instability, a key hall mark of cancer cells. Taken together, our data demonstrate we can control ERBB2 across many cancer types and as well control the major driver of cancers, which is aneuploidy without causing genome rearrangement. Future work will extend this technology to target more oncogenes across different disease settings and potentially any transcript of interest.

## Acknowledgement

We acknowledge the input, advice, and support of Dr. Kevin Struhl of the Harvard Medical School during the conception of this project. We thank Dr. Andrew Wolfe for cell lines provided, and Dr Frauke Stanke of the Hannover Medical School for her advice and support.

## Funding

National Cancer Institute (#U54CA221704 to Olorunseun O. Ogunwobi)

## Author’s contribution

Conceptualization: CUA, OOO

Methodology: CUA

Investigation: CUA, YG, FG, KY, LA, JZ, AA, FD, DW, OOO

Visualization: CUA, YG, FG, KY, LA, JZ, AA, FD, DW, OOO

Funding acquisition: OOO

Project administration: CUA, OOO

Supervision: OOO

Writing – original draft: CUA, OOO

Writing – review & editing: CUA, DW, OOO

CUA conceived the project and designed and engineered the destabilized 3’UTR AU rich elements and performed experiments. YG performed experiments. FL performed experiments, KY performed experiments, LA performed experiments, AA performed experiments. OOO conceived experiments in clinically relevant models and supervised experiments. JZ performed RNA Seq analysis. CUA, DW and OOO wrote the manuscript.

## Competing Interest

Chidiebere U Awah, Kevin Struhl and Olorunseun O Ogunwobi have filed for patent based on findings from this work.

## Data and material availability

Breast cancer cell line 3’UTR Sanger Sequencing, NCI H1975 **RNA** Seq data and Optical genome mapping data are deposited on Zenodo (10.5281/zenodo.6968947). All primary data and sequence and methods required to reproduce the conclusion of this paper is in the manuscript and supplementary data. CUNY and UTR Therapeutics Inc. will handle the requests for constructs created in this study.

---

**Supplementary Figure 1:**
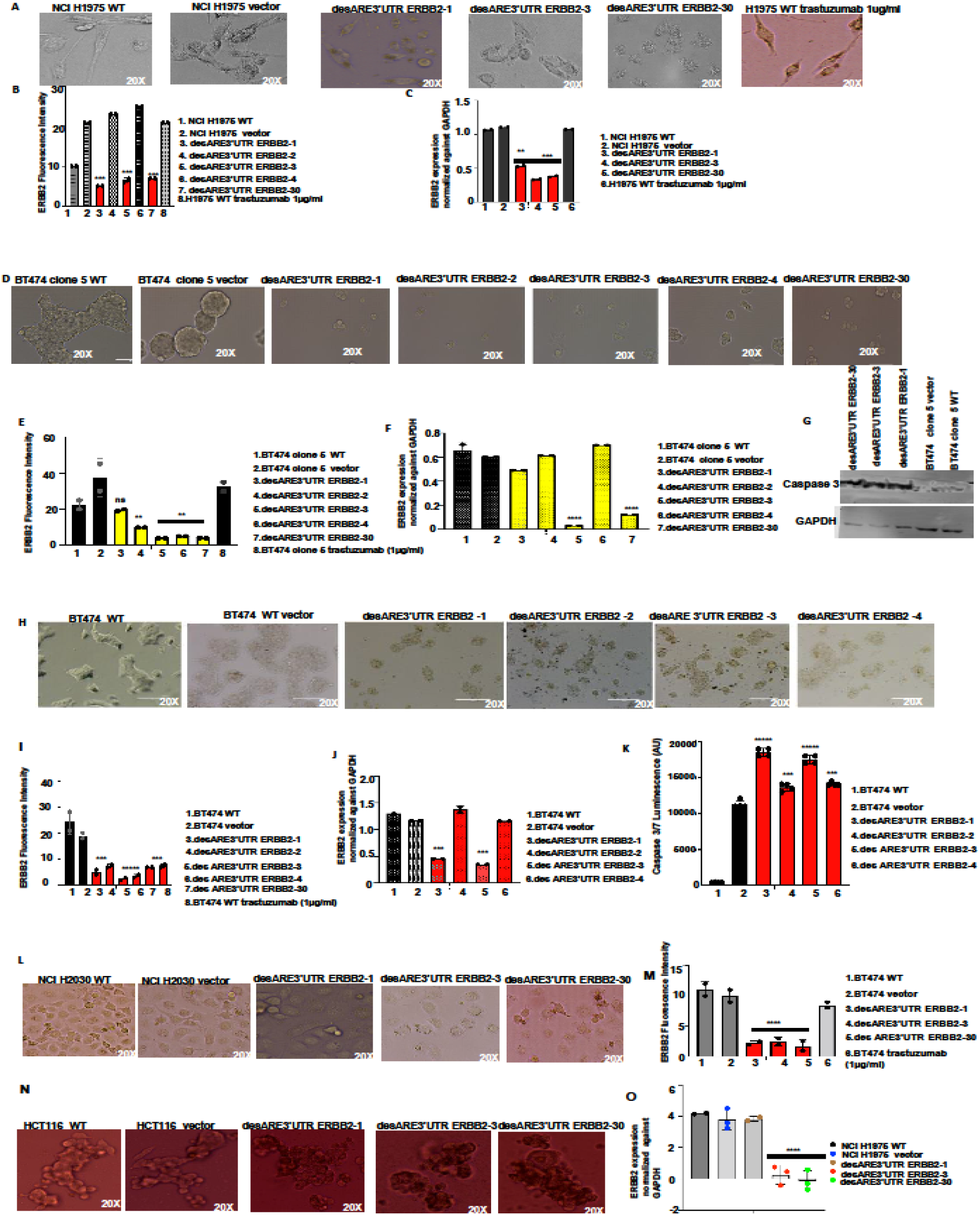
The destabilized 3’UTR ARE of ERBB2 severely distorted lung cancer and breast cancer cells membrane, degraded ERBB2 transcript and lead to cell death. **A.** Pictures show bright field microscope images of the NCI-H1975 wildtype cells, vector, cells containing constructs desARE3’UTR ERBB2-1, 3 and 30 and wild type trastuzumab treated NCI-H1975 cells (n=2). **B.** Bar charts show quantification of the ERBB2 fluorescence expression on NCI-H1975 wildtype cells, vector and cells containing constructs desARE3’UTR ERBB2-1, 3, 30 (red bars), desARE3’UTR ERBB2-2,4 and wild type trastuzumab treated NCI-H1975 cells (n=2). T-test *** =0.003, desARE3’UTRERBB2-1,3,30. **C.** Bar charts show quantification of the ERBB2 western blot expression normalized against GAPDH on NCI-H1975 wildtype cells, vector and cells containing constructs desARE3’UTR ERBB2-1, 3, 30 (red bars) and wild type trastuzumab treated NCI-H1975 cells (n=2). T-test *** =0.0021 desARE3’UTRERBB2-3,30, ** =0.02 desARE3’UTRERBB2-1 **D.** Pictures show bright field microscope images of the BT474 clone 5 wildtype cells, vector, cells containing constructs desARE3’UTR ERBB2-1,2, 3, 4 and 30 (n=3). **E.** Bar charts show quantification of the ERBB2 fluorescence expression on BT474 clone 5 wildtype cells, vector and cells containing constructs desARE3’UTR ERBB2-1,2, 3, 4, 30 and trastuzumab treated cells (yellow bars) (n=2). T-test, ns=desARE3’UTRERBB2-1, **=0.037 desARE3’UTRERBB2-2, **=0.0178 desARE3’UTRERBB2-3, **=0.0198 desARE3’UTRERBB2-4, ** =0.0178 desARE3’UTRERBB2-30. **F.** Bar charts show quantification of the ERBB2 western blot expression normalized against GAPDH on BT474 clone 5 wildtype cells, vector and cells containing constructs desARE3’UTR ERBB2-1, 2, 3, 4 and 30 (yellow bars) (n=3). T-test ****=0.00012 desARE3’UTRERBB2-3,30. **G.** Western blots show active caspase 3 and GAPDH expression across the wildtype BT474 clone 5, vector and on BT474 clone 5 containing constructs desARE3’UTR ERBB2-1, 3, and 30 (n=2). **H.** Pictures show bright field microscope images of the BT474 wildtype cells, vector, cells containing constructs desARE3’UTR ERBB2-1,2, 3, and 4 (n=3). **I.** Bar charts show quantification of the ERBB2 fluorescence expression on BT474 wildtype cells, vector and cells containing constructs desARE3’UTR ERBB2-1,2, 3, 4 (red bars) (n=2). T -test, **=0.0331 desARE3’UTRERBB2-1, **=0.0040 desARE3’UTRERBB2-2, **=0.0249 desARE3’UTRERBB2-3, **=0.0272 desARE3’UTRERBB2-4, **=0.0377 desARE3’UTRERBB2-30, **=0.0406 BT474 trastuzumab treated cells. **J.** Bar charts show quantification of the ERBB2 western blot expression normalized against GAPDH on BT474 wildtype cells, vector and cells containing constructs desARE3’UTR ERBB2-1, 2, 3, and 4 (red bars) (n=2). T-test, ***=0.0036 desARE3’UTRERBB2-1 and 3. **K.** Bar charts show active caspase 3/7 luminescence on BT474 wildtype cells, vector and cells containing constructs desARE3’UTR ERBB2-1, 2, 3 and 4 (n=2). T-test, *****<0.00001 desARE3’UTRERBB2-1,3. ***=0.0022 desARE3’UTRERBB2-2,4. **L.** Pictures show bright field microscope images of the NCI-H2030 wildtype cells, vector, cells containing constructs desARE3’UTR ERBB2-1, 3 and 30 (n=2). **M.** Bar charts show quantification of the ERBB2 fluorescence intensity on NCI H2030 wildtype cells, vector and cells containing constructs desARE3’UTR ERBB2-1, 3, 30 and trastuzumab treated cells (n=2). T-test, ****=0.0004 desARE3’UTRERBB2-1,3,30. **N.** Pictures show bright field microscope images of the HCT116 wildtype cells, vector, cells containing constructs desARE3’UTR ERBB2-1, 3 and 30 (n=2). **O.** Bar charts show quantification of the ERBB2 expression normalized against GAPDH by qPCR on HCT116 wildtype cells, vector and HCT116 cells containing constructs desARE3’UTR ERBB2-1, 3 and 30 (n=2). T-test (****=0.0003, desARE3’UTRERBB2-3 and 30).

**Supplementary Figure 2:**
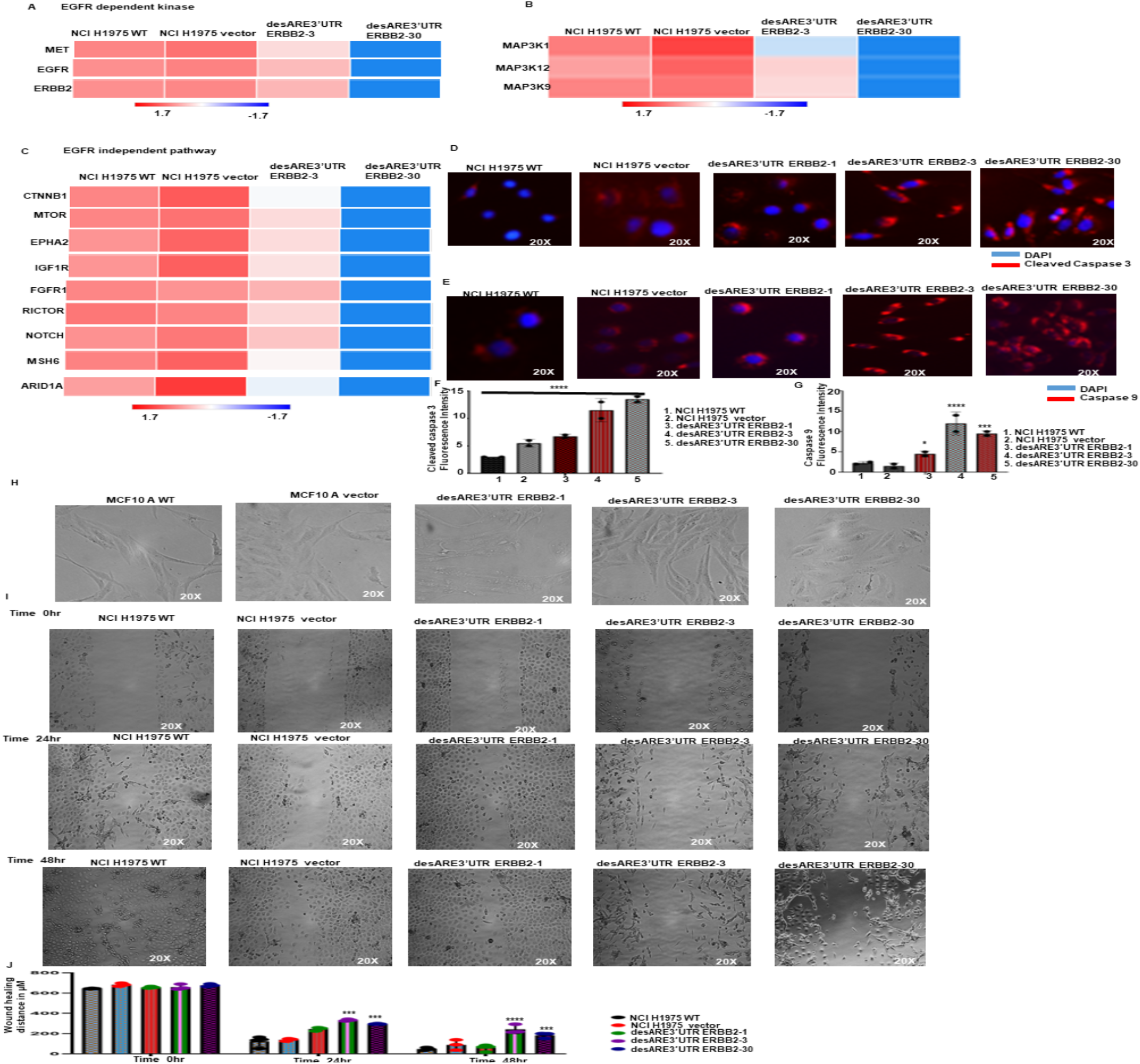
Engineered destabilized 3’UTR ARE of ERBB2 degradation of ERBB2 in EGFR T90M reduces the expression of EGFR dependent and independent kinases in non-small cell lung cancer. **A.** Heat map shows EGFR dependent kinases known to cause osimertinib resistance: EGFR, MET and ERBB2 expression changes in the wildtype NCI H1975, vector, desARE3’UTR ERBB2-3 and 30. **B.** Heat map shows EGFR dependent kinases known to cause osimertinib resistance: MAP3K1, MAP3K12, MAP3K9 expression changes in the wildtype NCI H1975, vector, desARE3’UTR ERBB2-3 and 30. **C.** Heat map shows EGFR independent genes known to cause osimertinib drug resistance: CTNNB1, MTOR, EPHA2, IGFR1, FGFR1, RICTOR, NOTCH, MSH6, ARID1A expression changes in the wildtype NCI H1975, vector, desARE3’UTR ERBB2-3 and 30. **D.** Immunofluorescence images show cleaved caspase 3 expression (stained in red) and nuclei (DAPI-blue) on wildtype NCI H1975, vector and on NCI H1975 containing constructs desARE3’UTR ERBB2-1, 3, 30 (n=2). **E.** Immunofluorescence images show caspase 9 expression (stained in red) and nuclei (DAPI-blue) on wildtype NCI H1975, vector and on NCI H1975 containing constructs desARE3’UTR ERBB2-1, 3, 30 (n=2). **F.** Bar charts show quantification of the cleaved caspase 3 fluorescence intensity on NCI H1975 wildtype cells, vector and cells containing constructs desARE3’UTR ERBB2-1, 3, 30 (n=2). T-test, ****=0.00025 desARE3’UTRERBB2-3,30 **G.** Bar charts show quantification of the caspase 9 fluorescence intensity on NCI H1975 wildtype cells, vector and cells containing constructs desARE3’UTR ERBB2-1, 3, 30 (n=2). T-test, * p=0.05 desARE3’UTR ERBB2-1, ****p=0.00025 desARE3’UTRERBB2-3,30 ***p=0.0022 desARE3’UTR ERBB2-30. **H.** Bright field microscope images of the MCF10A wildtype cells, vector, cells containing constructs desARE3’UTR ERBB2-1, 3 and 30. **I.** Bright field microscope images of wound scratch and closing in wild type NCI H1975, vector, desARE3’UTR ERBB2-1,3,30 treated cells at 0hr, 22hr and 48hr. **J.** Bar chart shows the quantification of wound opening at 0hr, 22 and 48hrs in NCI H1975 wildtype, vector, desARE3’UTR1,3,30. At 22hr desARE3’UTRERBB2-3,30 show significant wound opening t-test ***p=0.0014 and at 48hr t-test **** p=0.0004 desARE3’UTRERBB2-3, ***p=0.003 desARE3’UTRERBB2-30. All experiment n=2.

**Supplementary Figure 3:**
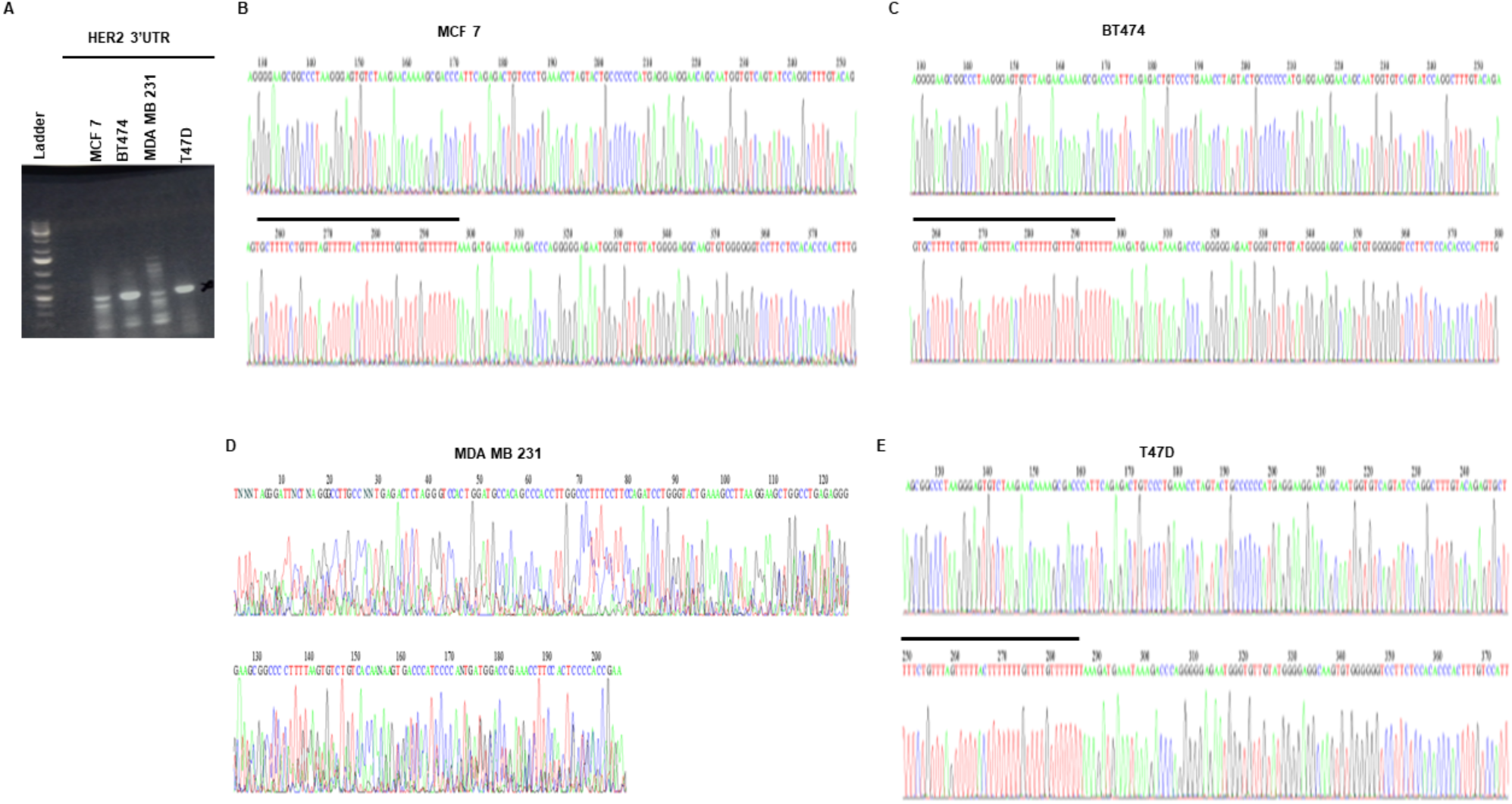
Identification of poly U stabilizing sequences on the 3’UTR of ERBB2. **A.** Gel picture shows the cDNA amplification of ERBB2 3’UTR across different cancers band size is 450bp marked in black asterisks. **B.** Sanger sequencing electropherogram of ERBB2 3’UTR cDNA. The poly U stabilizing sequences are marked in black line in MCF7. **C.** Sanger sequencing electropherogram of ERBB2 3’UTR cDNA. The poly U stabilizing sequences are marked in black line in BT474. **D.** Sanger sequencing electropherogram of ERBB2 3’UTR cDNA. The poly U stabilizing sequences are not present in the triple negative breast cancer MDAMB231 **E.** Sanger sequencing electropherogram of ERBB2 3’UTR cDNA. The poly U stabilizing sequences are marked in black line in T47D.

**Supplementary Figure 4:**
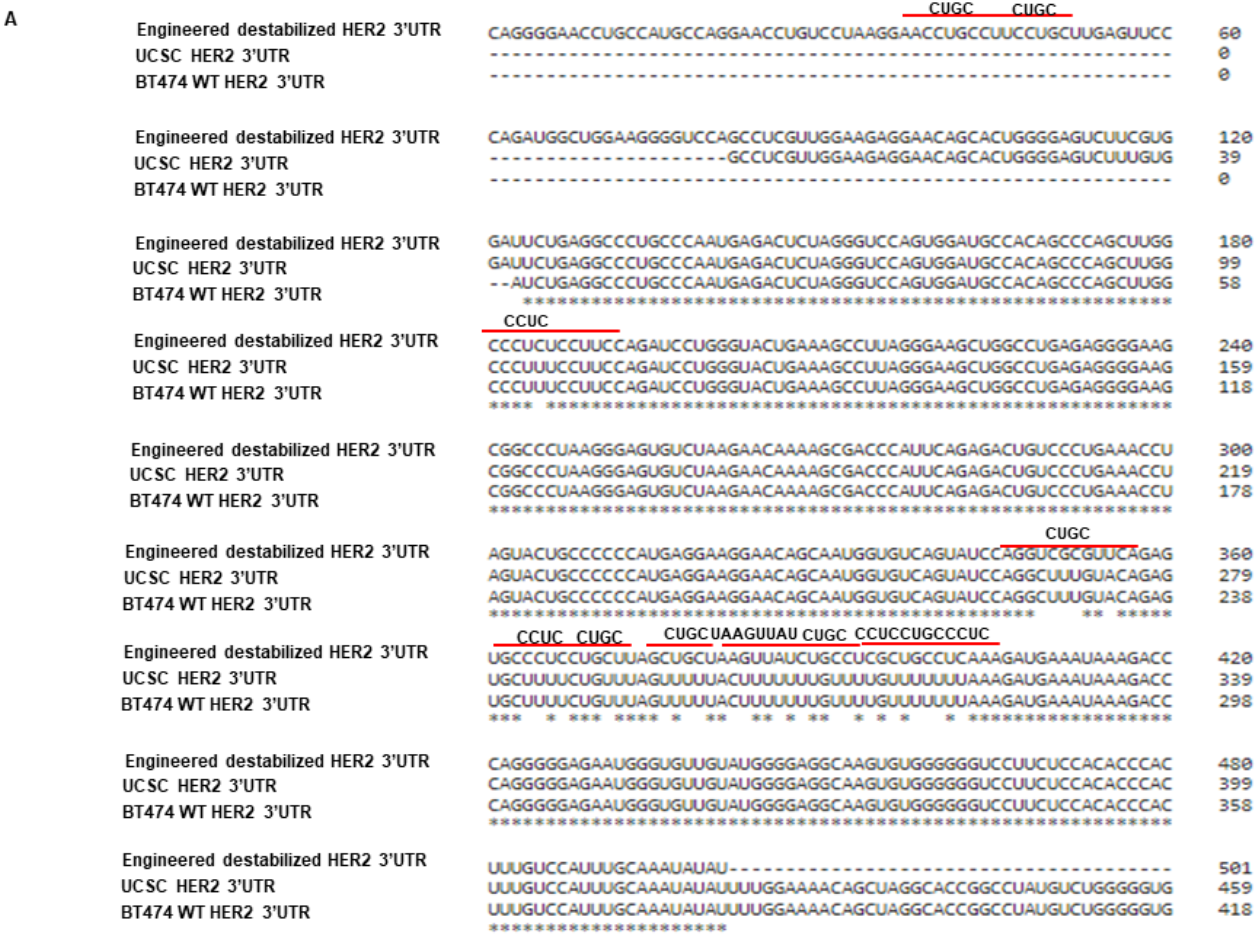
ERBB2 3’UTR cDNA converted to mRNA sequences with stabilizing poly U engineered to destabilizing elements underlined in red. **A.** Sequence alignment of the HER 3’UTR sequence from the UCSC genome browser, ERBB2 3’UTR sequence from the BT474 WT and the engineered ERBB2 3’UTR sequence. The sequence underlined in red line are the motif we replaced the stabilizing elements with.

**Supplementary Figure 5:**
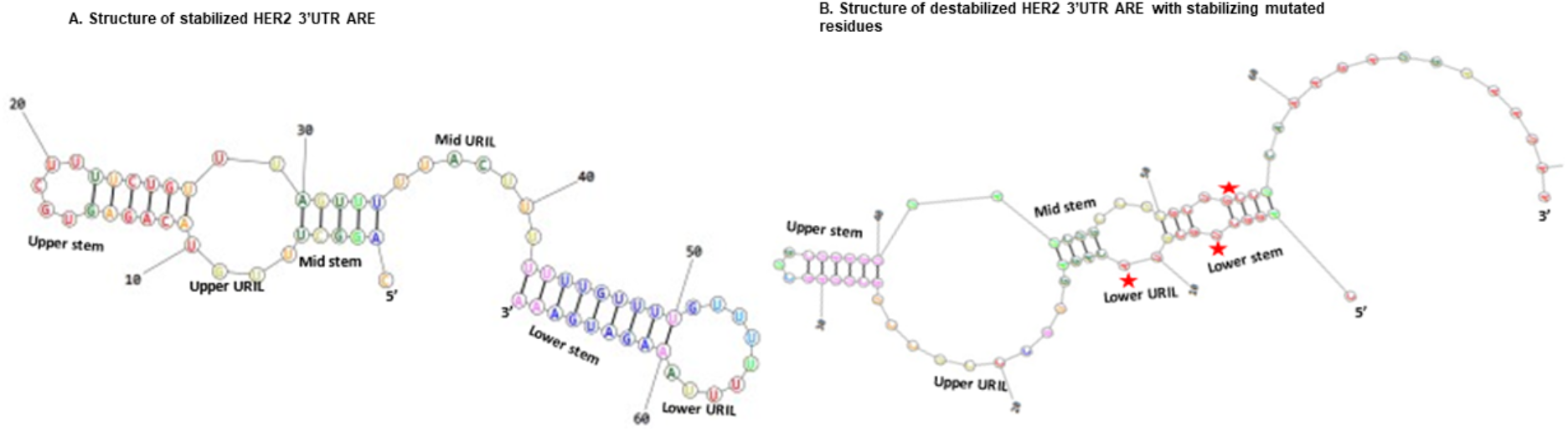
Structure model of stable ERBB2 3’UTR ARE and destabilized ERBB2 3’UTR ARE motif with increased stability due to mutated residues marked in red. **A-B.** Depiction of component of the stabilized and destabilized 3’UTR ARE, asterisk in **B**represents residues that will mutated to enhance the stability of destabilized 3’UTR ARE elements.

**Supplementary Figure 6:**
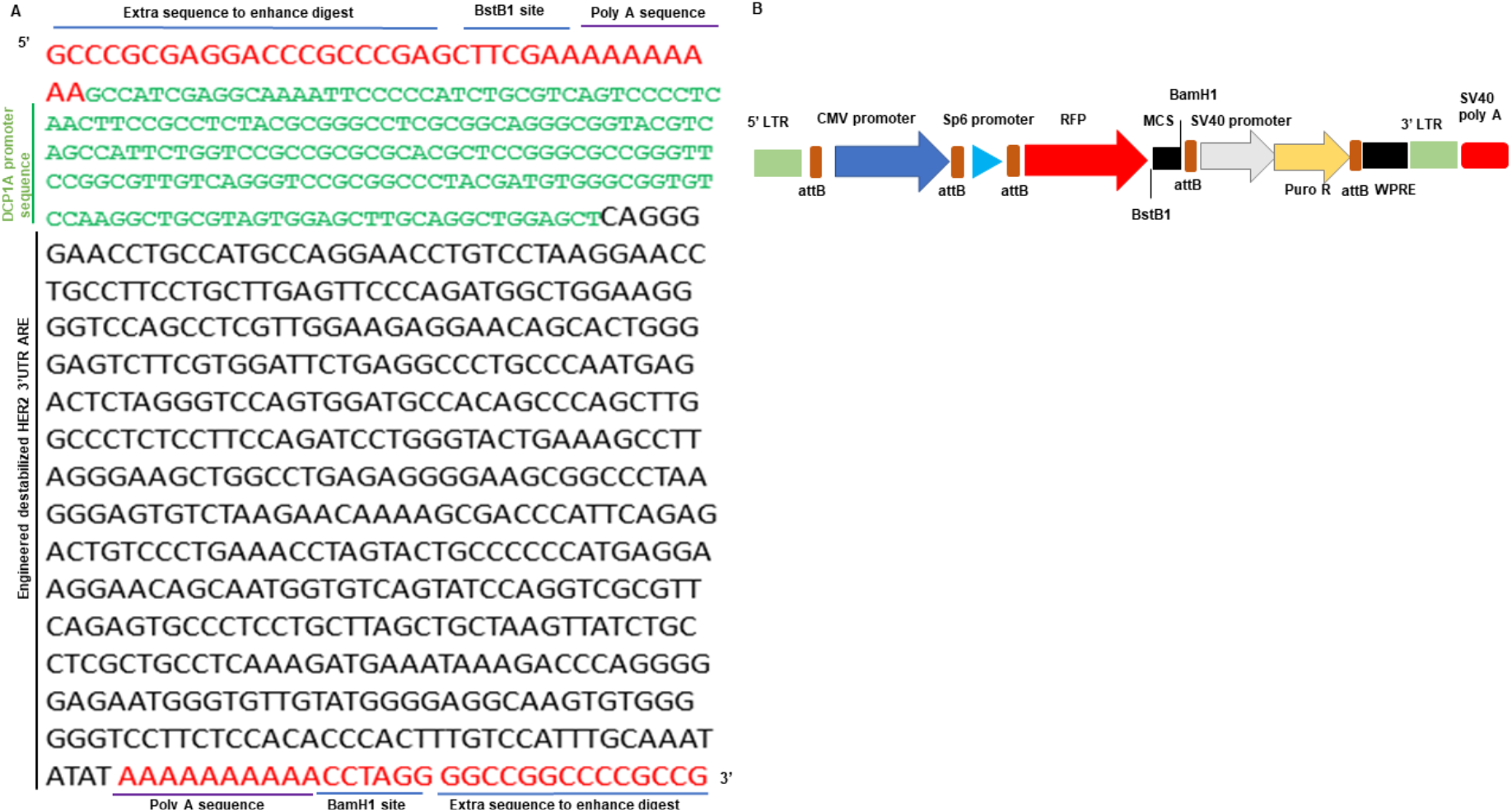
Design of destabilized ERBB2 3’UTR ARE with DCP1A promoter, BstB1, BamH1 restriction sites and plasmid vector design. **A.** Schematic depiction of the design of the synthetic gblock showing nucleotides to enhance restriction digest, BstB1 sites, Poly A sequence, hDCP1A promoter (green), the engineered destabilized ERBB2 3’UTR ARE (black) and BamH1 sites. **B.** Representation of the structure of the vector depicting the components parts and the restriction enzyme sites used at the multiple cloning site.

**Supplementary Figure 7:**
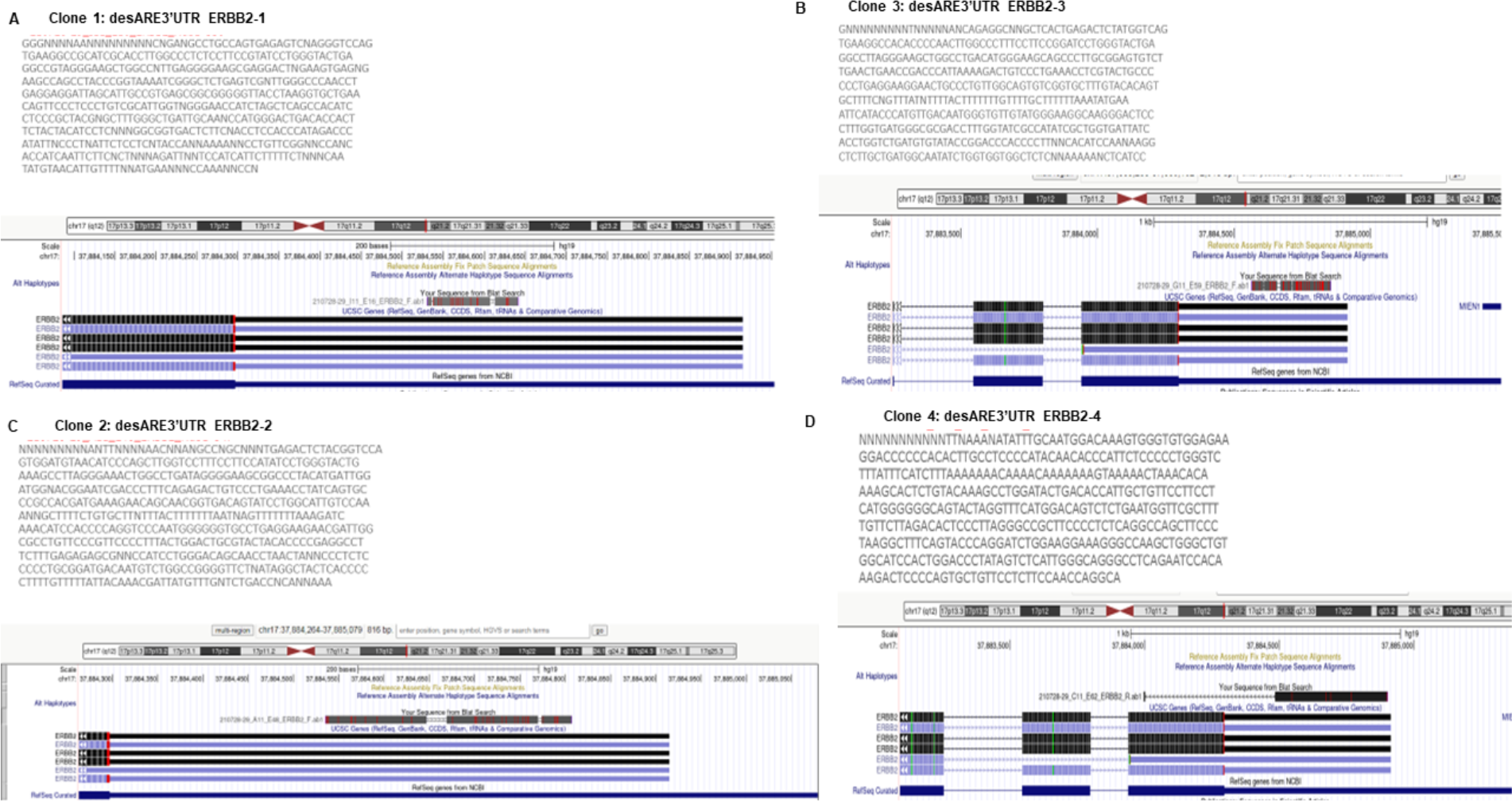
Identification of successful engineered destabilized ERBB2 3’UTR ARE obtained by cloning in recombinant efficient E. coli. **A-D.** Sequences of desARE3’UTR ERBB2-1, 2,3 and 4 clones mapped to the ERBB2 3’UTR

**Supplementary Figure 8:**
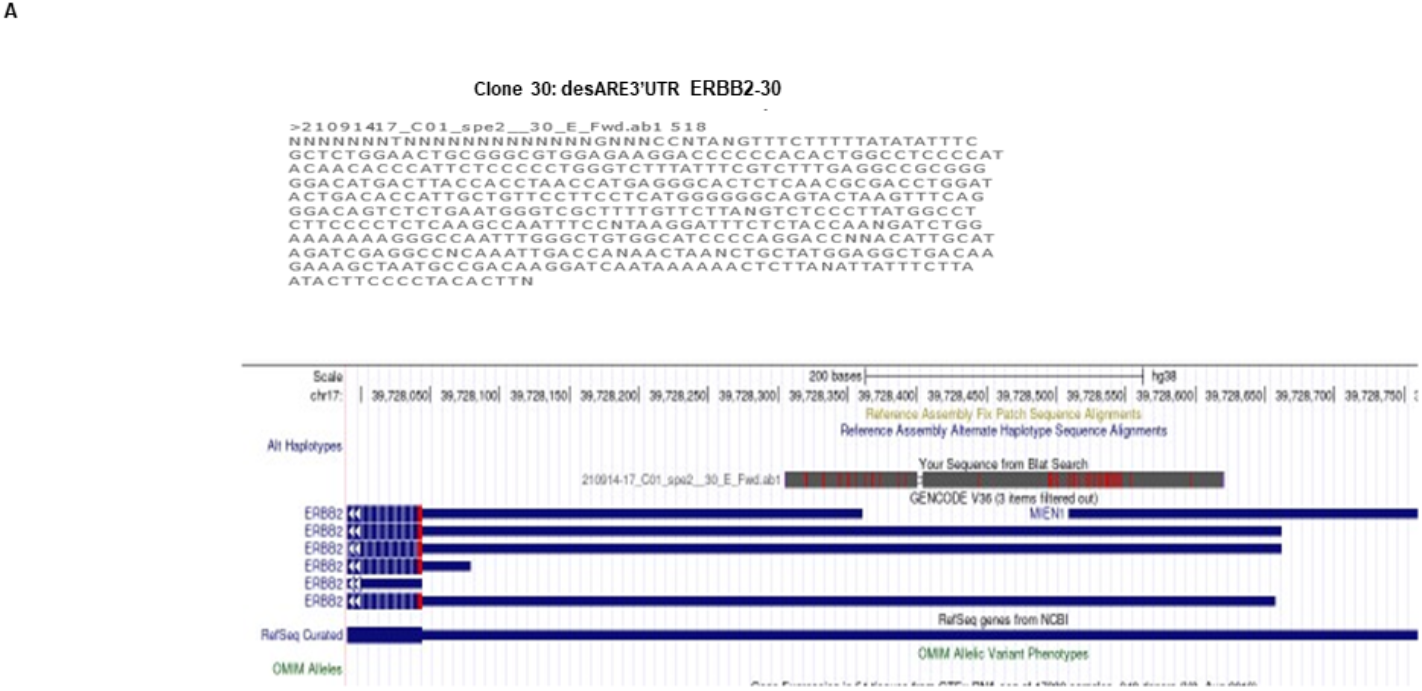
Identification of successful engineered destabilized ERBB2 3’UTR ARE obtained by Gibson assembly cloned in recombinant deficient E. coli. **A.** Sequence of desARE3’UTR ERBB2-30 clone mapped to the ERBB2 3’UTR.

**Supplementary Figure 9:**
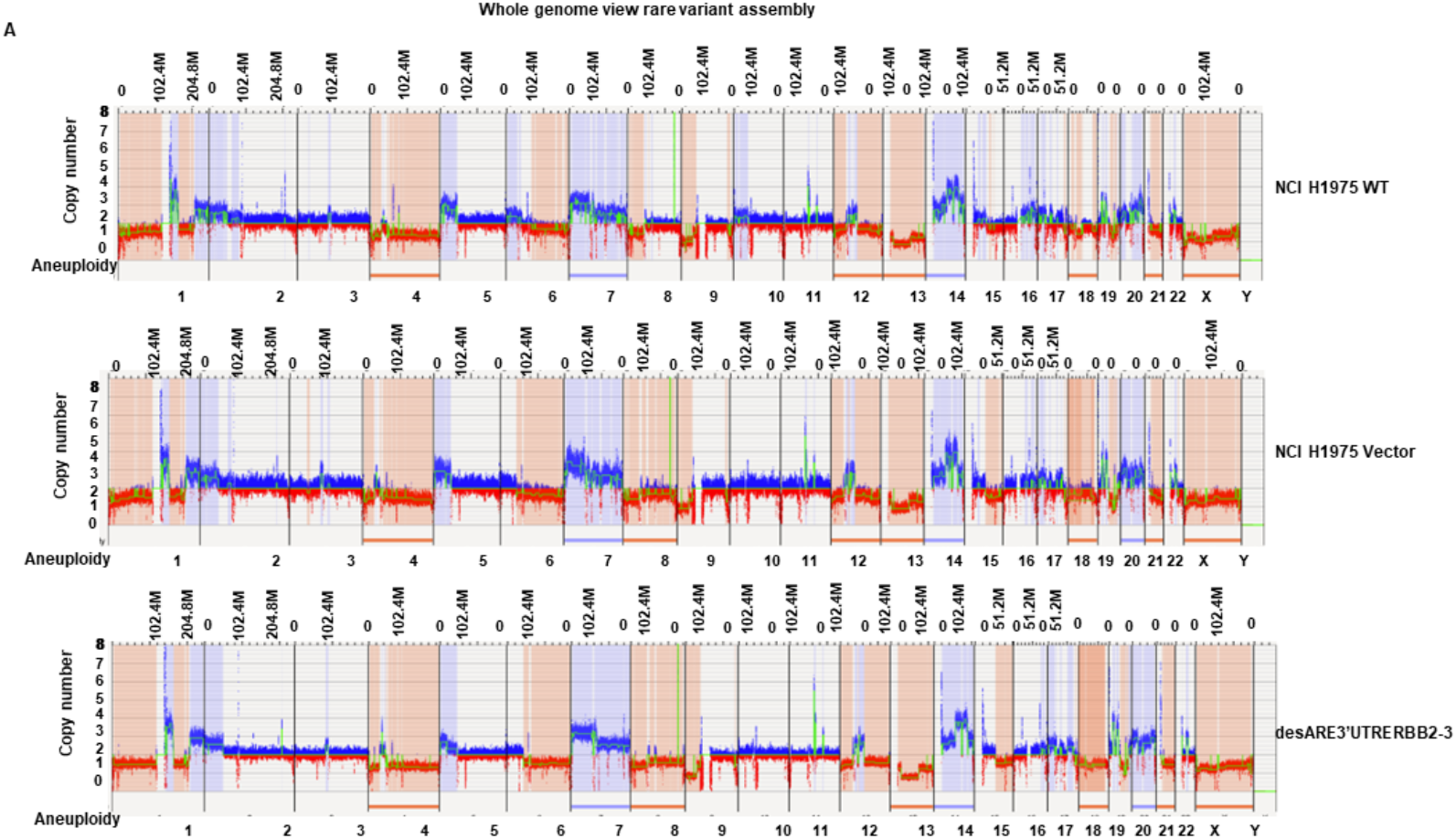
Whole genome view of rare variant assembly comparing wild type EGFR T790M, vector and desARE3’UTR ERBB2-3 non-small cell lung cancer cells. **A.** Genome view of whole rare variant assembly in wild type NCI H1975, vector and desARE3’UTRERBB2-3 n=2

**Supplementary Figure 10:**
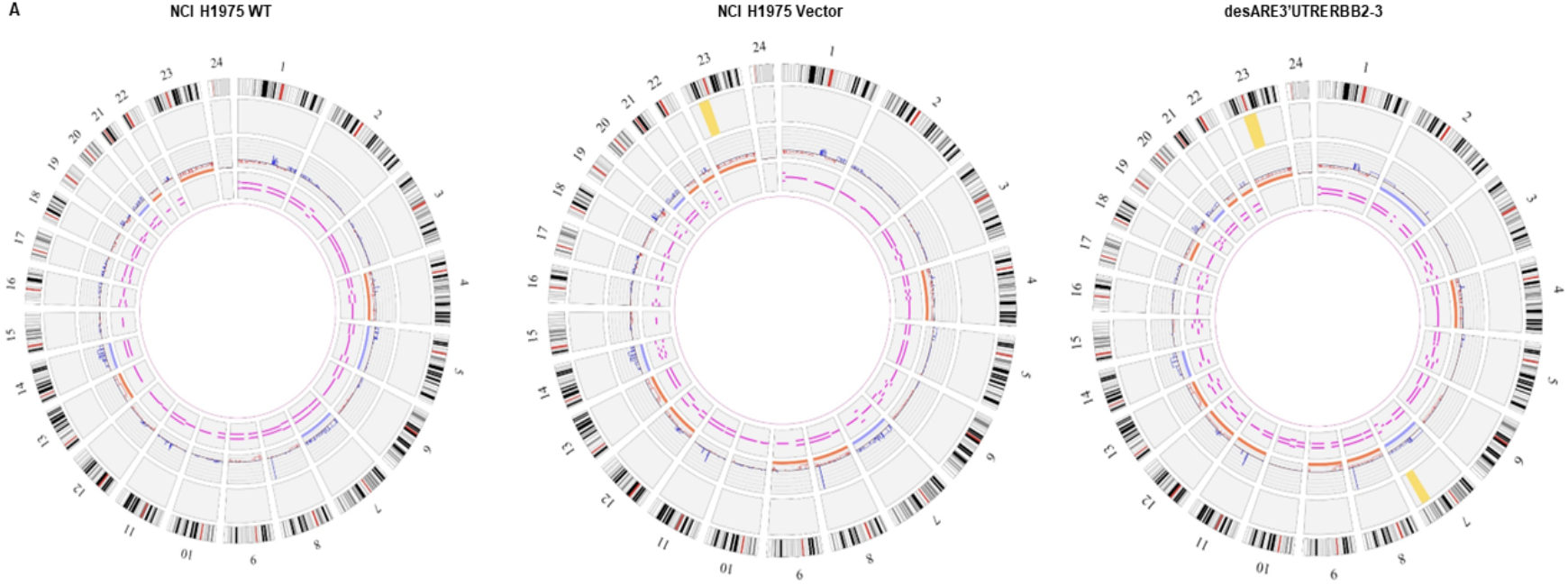
Whole genome view of rare variant assembly comparing wild type EGFR T790M, vector and desARE3’UTR ERBB2-3 non-small cell lung cancer cells. **A.** Circos plot of de novo assembly in wild type NCI H1975, vector and desARE3’UTRERBB2-3. n=2

**Supplementary Figure 11:**
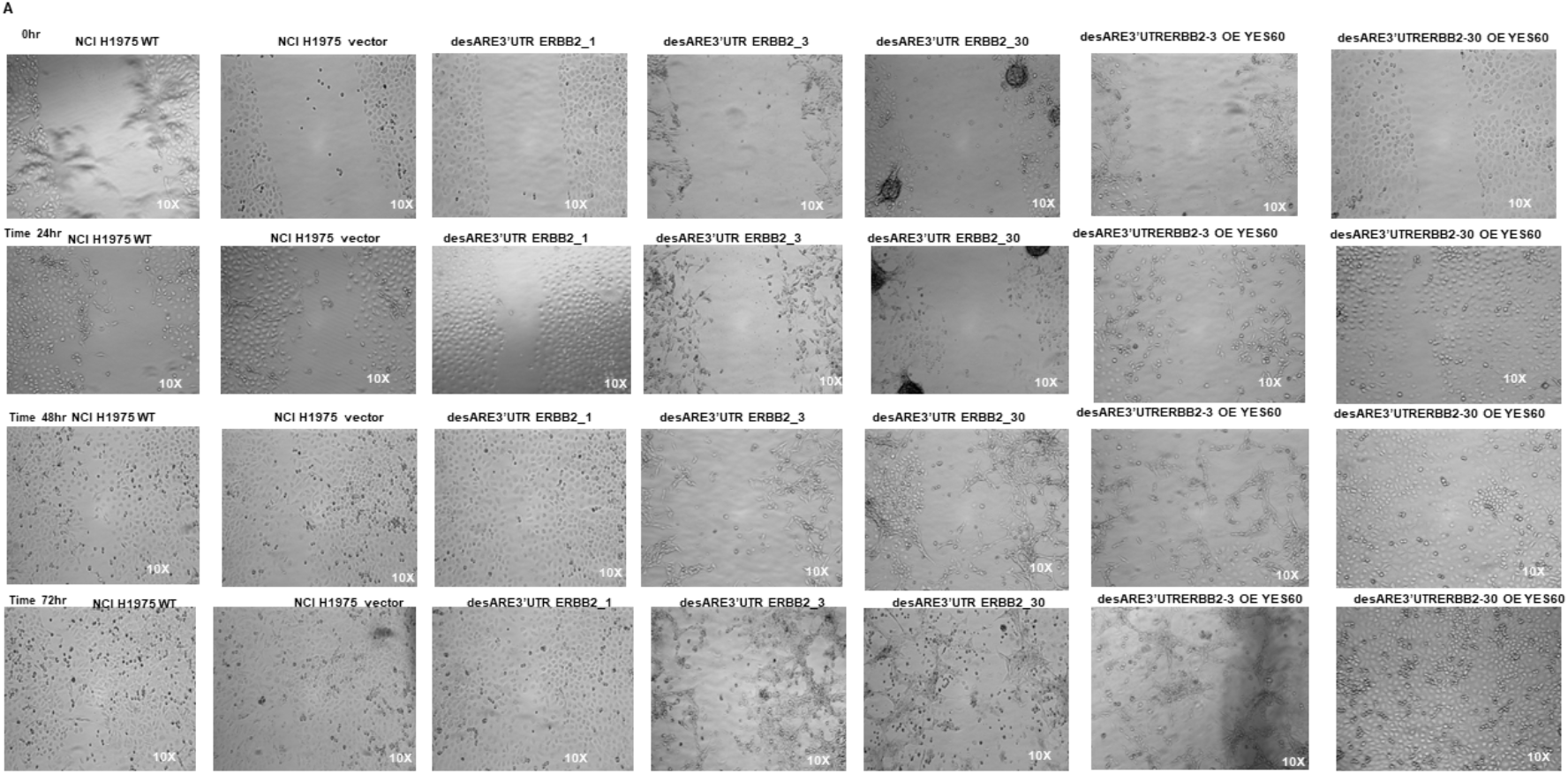
Migration assay of NCI H1975 WT, vector, desARE 3’UTRERBB2-1, 3, 30 and destabilized cells with over expressed YES1. **A.** Microscope images of wound healing assay of NCI H1975 WT, vector, desARE3’UTRERBB2-1, 3 and 30 and desARE3’UTRERBB2-3 and 30 overexpressed with YES1.Images were taken at 0hr, 24hr, 48hrs and 72hrs at 10x.

**Supplementary Figure 12:**
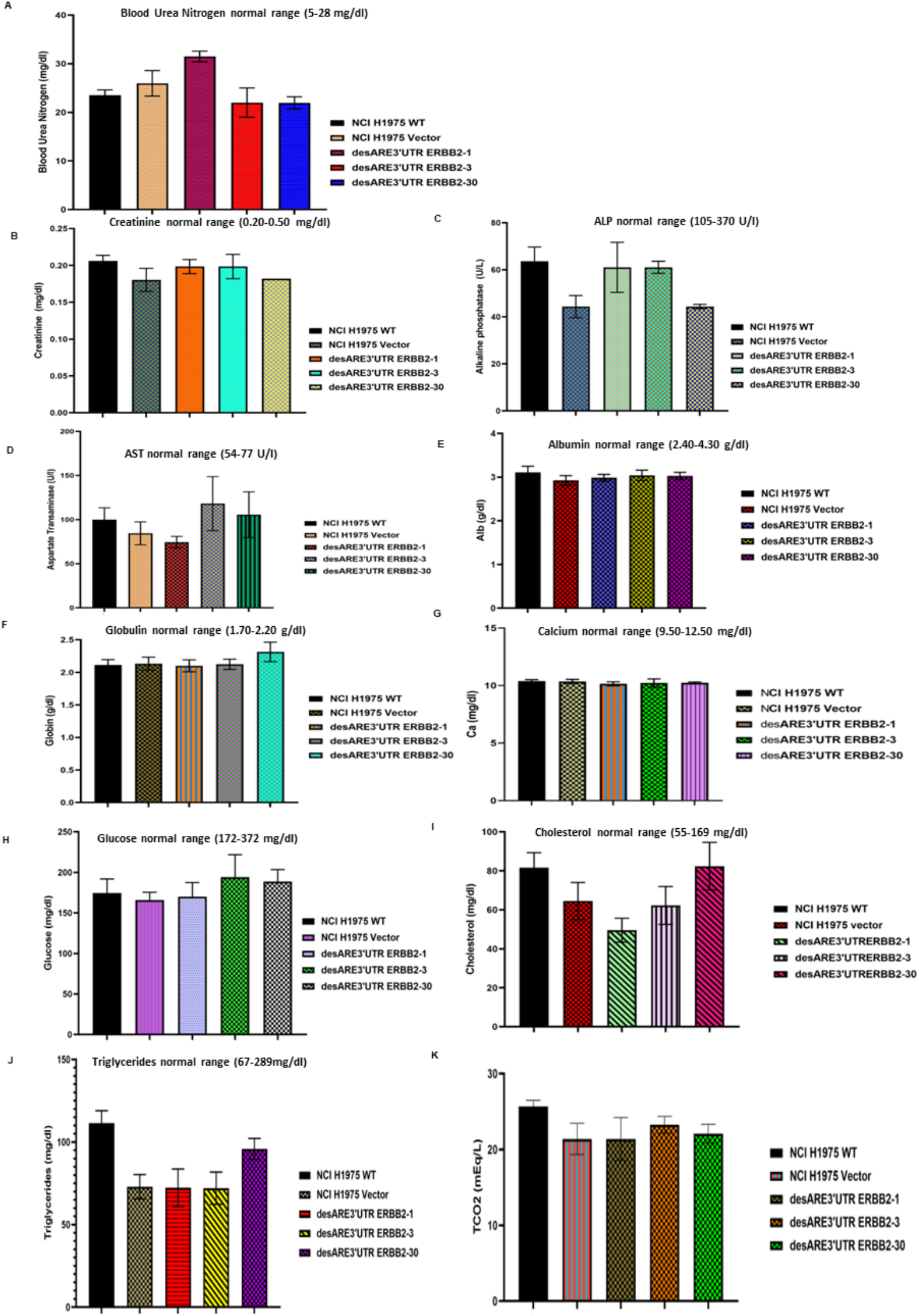
Engineered destabilized ARE 3’UTR of ERBB2 does not affect the vital organ function not normal cell electrolyte balance. **A.** Bar charts show blood urea nitrogen level of mice bearing tumor NCI H1975 WT (black), vector (yellow), and mice treated with desARE3’UTRERBB2-1(light green), desARE3’UTRERBB2-3 (green) and desARE3’UTRERBB2-30 (red). **B.** Bar charts show creatinine levels of mice bearing tumor NCI H1975 WT (black), vector (gray), and mice treated with desARE3’UTRERBB2-1(orange), desARE3’UTRERBB2-3 (light green) and desARE3’UTRERBB2-30 (brown). **C.** Bar charts show alkaline phosphatase level of mice bearing tumor NCI H1975 WT (black), vector (blue), and mice treated with desARE3’UTRERBB2-1(light green), desARE3’UTRERBB2-3 (dark green) and desARE3’UTRERBB2-30 (white on black crosses). **D.** Bar charts show aspartate amino transferase of mice bearing tumor NCI H1975 WT (black), vector (orange), and mice treated with desARE3’UTRERBB2-1(red), desARE3’UTRERBB2-3 (white on black crosses) and desARE3’UTRERBB2-30 (green with black lines). **E.** Bar charts show albumin level of mice bearing tumor NCI H1975 WT (black), vector (red), and mice treated with desARE3’UTRERBB2-1(blue), desARE3’UTRERBB2-3 (orange) and desARE3’UTRERBB2-30 (purple). **F.** Bar charts show globulin of mice bearing tumor NCI H1975 WT (black), vector (yellow stripped), and mice treated with desARE3’UTRERBB2-1(orange), desARE3’UTRERBB2-3 (white shaded) and desARE3’UTRERBB2-30 (light green). **G.** Bar charts show calcium level of mice bearing tumor NCI H1975 WT (black), vector (yellow stripped), and mice treated with desARE3’UTRERBB2-1(blue stripped), desARE3’UTRERBB2-3 (green stripped) and desARE3’UTRERBB2-30 (pink stripped). **H.** Bar charts show glucose level of mice bearing tumor NCI H1975 WT (black), vector (pink), and mice treated with desARE3’UTRERBB2-1(light blue), desARE3’UTRERBB2-3 (green stripped) and desARE3’UTRERBB2-30 (black stripes on white). **I.** Bar charts show cholesterol level of mice bearing tumor NCI H1975 WT (black), vector (red), and mice treated with desARE3’UTRERBB2-1(green stripe), desARE3’UTRERBB2-3 (light pink) and desARE3’UTRERBB2-30 (purple stripe). **J.** Bar charts show triglyceride level of mice bearing tumor NCI H1975 WT (black), vector (light yellow), and mice treated with desARE3’UTRERBB2-1(red stripes), desARE3’UTRERBB2-3 (bright yellow stripes) and desARE3’UTRERBB2-30 (purple stripes). **K.** Bar charts show blood level of C02 of mice bearing tumor NCI H1975 WT (black), vector (red stripes), and mice treated with desARE3’UTRERBB2-1(orange stripes), desARE3’UTRERBB2-3 (yellow stripes) and desARE3’UTRERBB2-30 (green stripes).

**Supplementary Figure 13:**
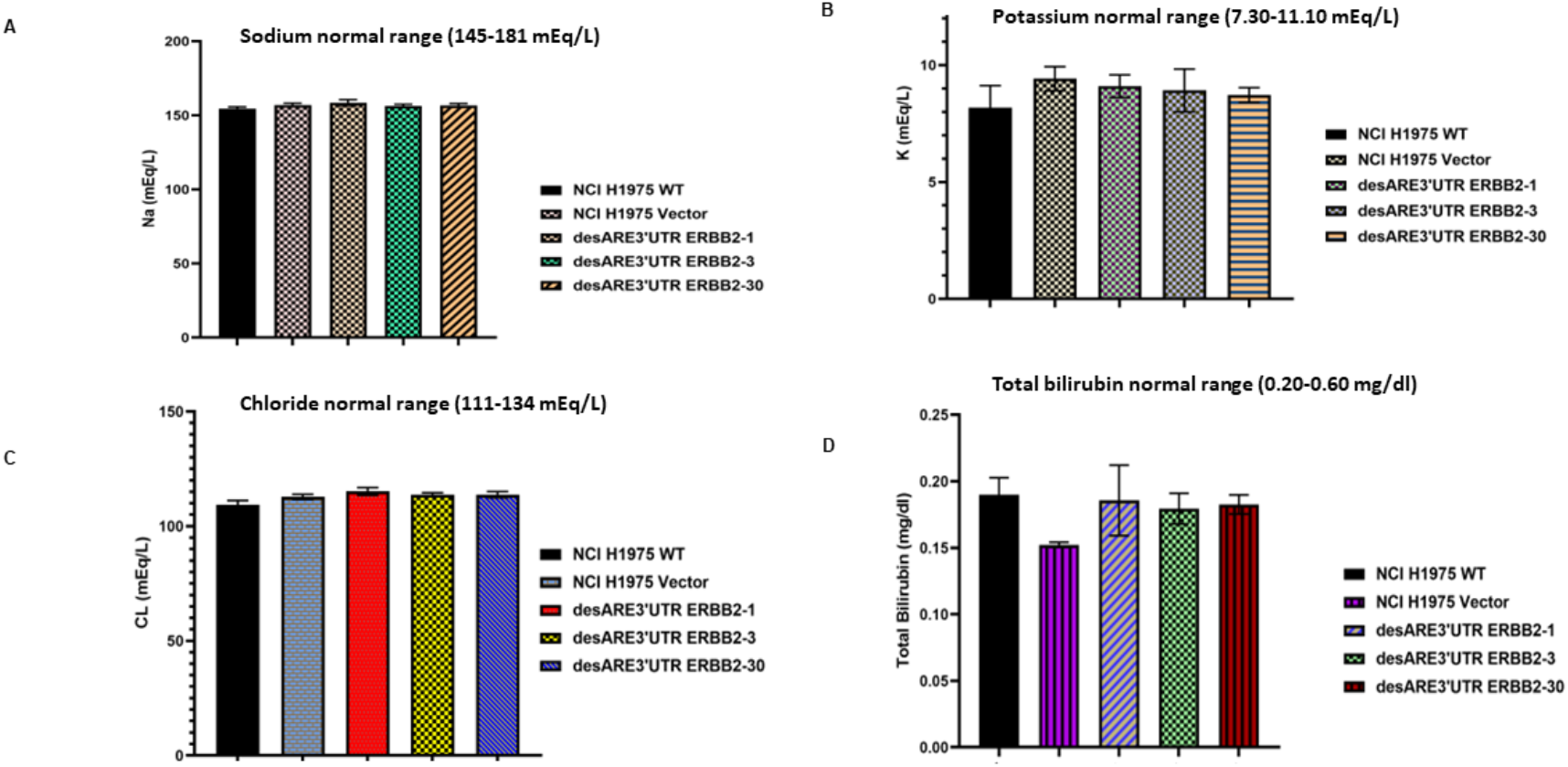
Analysis of electrolyte levels in NCI H1975 WT, vector in mice bearing tumors and in mice bearing tumors treated with desARE3’UTR ERBB2-1,3 and 30. **A.** Bar charts show blood sodium level of mice bearing tumor NCI H1975 WT (black), vector (pink), and mice treated with desARE3’UTRERBB2-1(light blue stripes), desARE3’UTRERBB2-3 (green stripes) and desARE3’UTRERBB2-30 (yellow stripes). **B.** Bar charts show blood potassium level of mice bearing tumor NCI H1975 WT (black), vector (yellow stripes), and mice treated with desARE3’UTRERBB2-1(green stripes), desARE3’UTRERBB2-3 (blue stripes) and desARE3’UTRERBB2-30 (yellow stripes) **C.** Bar charts show blood chloride of mice bearing tumor NCI H1975 WT (black), vector (blue), and mice treated with desARE3’UTRERBB2-1(red), desARE3’UTRERBB2-3 (yellow) and desARE3’UTRERBB2-30 (blue stripes). **D.** Bar charts show total bilirubin levels of mice bearing tumor NCI H1975 WT (black), vector (pink), and mice treated with desARE3’UTRERBB2-1(blue), desARE3’UTRERBB2-3 (green stripes) and desARE3’UTRERBB2-30 (red stripes).

**Supplementary Figure 14.**
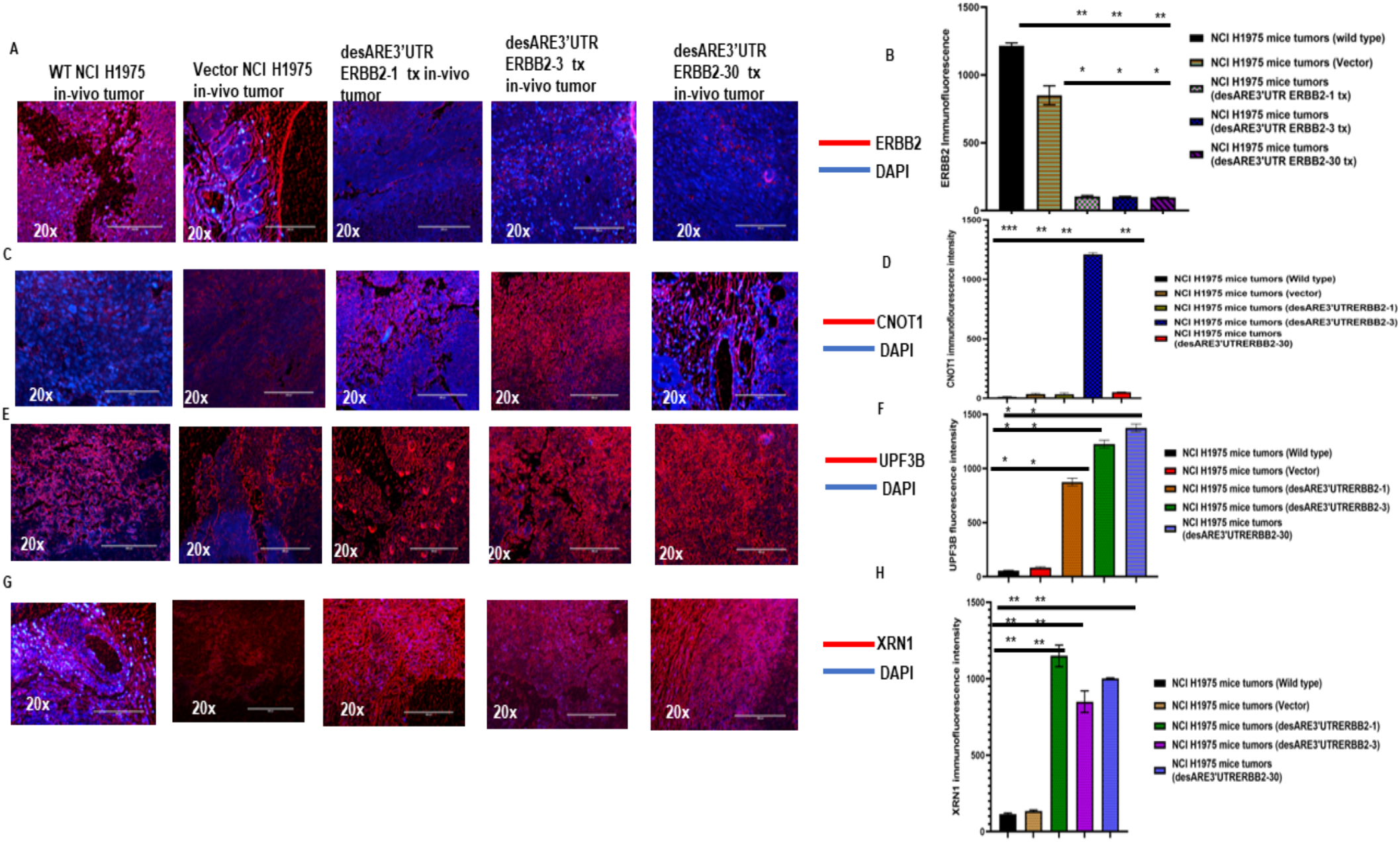
Immunofluorescence staining of xenografts tumors for ERBB2, CNOT1, UPF3B and XRN1. **A.** Immunofluorescence pictures show ERBB2 expression (stained in red) and nuclei (DAPI-blue) on xenograft tumors, wildtype NCI-H1975, vector and on tumors treated with constructs desARE3’UTR ERBB2-1, 3 and 30. **B.** Bar chart show quantification of ERBB2 protein fluorescence intensity on xenograft tumors, wildtype NCI-H1975, vector and on tumors treated with constructs desARE3’UTR ERBB2-1, 3 and 30. T-test of WT vs desARE3’UTR ERBB2-1,3,30 (p=0.0074, 0.0057, 0.0094). T-test on Vector vs desARE3’UTRERBB2-1,3,30 (p=0.038, 0.0409, 0.0434) **C.** Immunofluorescence pictures show CNOT1 expression (stained in red) and nuclei (DAPI-blue) on xenograft tumors, wildtype NCI-H1975, vector and on tumors treated with constructs desARE3’UTR ERBB2-1, 3 and 30. **D.** Bar chart show quantification of CNOT1 protein fluorescence intensity on xenograft tumors, wildtype NCI-H1975, vector and on tumors treated with constructs desARE3’UTR ERBB2-1, 3 and 30. T-T-test of desARE3’UTR ERBB2-3 vs WT, vector, desARE3’UTR ERBB2-1,30 (p=0.0040, 0.0008, 0.0027, 0.0163). **E.** Immunofluorescence pictures show UPF3B expression (stained in red) and nuclei (DAPI-blue) on xenograft tumors, wildtype NCI-H1975, vector and on tumors treated with constructs desARE3’UTR ERBB2-1, 3 and 30. **F.** Bar chart show quantification of UPF3B protein fluorescence intensity on xenograft tumors, wildtype NCI-H1975, vector and on tumors treated with constructs desARE3’UTR ERBB2-1, 3 and 30. T-test of WT vs desARE3’UTRERBB2-1,3,30 (p=0.023, 0.019, 0.0145). T-test of Vector vs desARE3’UTR ERBB2-1,3,30 (p=0.0242, 0.0112, 0.0148) **G.** Immunofluorescence pictures show XRN1 expression (stained in red) and nuclei (DAPI-blue) on xenograft tumors, wildtype NCI-H1975, vector and on tumors treated with constructs desARE3’UTR ERBB2-1, 3 and 30. **H.** Bar chart show quantification of UPF3B protein fluorescence intensity on xenograft tumors, wildtype NCI-H1975, vector and on tumors treated with constructs desARE3’UTR ERBB2-1, 3 and 30. T-test of WT vs desARE3’UTRERBB2-1,3,30 (p=0.027, 0.0475, 0.0054). T-test on Vector vs desARE3’UTR ERBB2-1,3,30 (p=0.028, 0.0489, 0.0055)

## Supplementary Materials

### Materials and Methods

#### Cell culture of BT474, BT474 clone 5, NCI H1975, NCI H2030, HCT116 and MCF10A

BT474 was grown in DMEM supplemented with 10% FBS and 1% penicillin-streptomycin. The BT474 clone 5 (trastuzumab resistant) and NCI H1975 EGFR T790M and NCI H2030 were grown in RPMI-1640 supplemented with 10% FBS and 1% penicillin-streptomycin. HCT116 were grown in McCoy’s 5A modified medium with 10% FBS and 1% penicillin-streptomycin. MCF10A were grown in MEBM with additives as listed in ATCC. Cells were grown to 80% confluency before use.

#### RNA extraction from breast cancer cell lines

We extracted total RNA from the BT474 using the RNA Easy kit from the Qiagen (cat no: 74104). The RNA was stored at -80C until use.

#### QPCR with ERBB2 3’UTR primers

To determine the stabilizing AU rich elements on the 3’UTR of ERBB2, we made cDNA from the total RNA of BT474, MCF7, T47D and MDA MB231 using the Qiagen reverse transcription kit (Catalog no: 205311). We performed RT-PCR using the cDNA according to the Qiagen manufacturer’s protocol using the following primer sequences:

ERBB2 forward: GCCTCGTTGGAAGAGGAACA

ERBB2 reverse: AGAGCCACCCCCAGACATAG

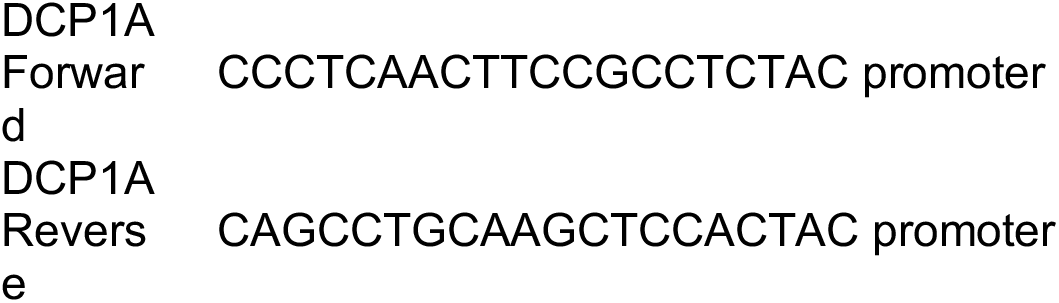

By using the following PCR cycle:

Expected ERBB2 3’UTR amplicon size = 464bp

PCR Cycle for cDNA synthesis:

94°C for 2mins

94°C for 15sec

55°C for 30sec

68-72 for 1min

Step2-3 for 40 cycles

4°C hold

PCR set up

38.1µl of H20

1.5µl of 50mM MgCl_2_

1µl of 10mM DNTP

1µl ERBB2 Forward primers

1µl ERBB2 Reverse primers

2µl of cDNA (200ng).

1µl of Taq polymerase

The amplicon was separated on 2% agarose gel.

#### Sanger sequencing of 3’UTR of ERBB2 cDNA amplicon

To confirm the ERBB2 stabilizing 3’UTR ARE. The amplified bands were excised under UV light and then extracted with Qiagen gel extraction kit (Catalog no: 28706X4) according to manufacturer’s protocol. The amplicon was then sequenced with the ERBB2 3’UTR PCR primers by Sanger Sequencing method by the commercial company Psomagen Inc. Brooklyn New York USA (**Fig. S3**). Subsequently, we used the online software DNA to mRNA translator (http://biomodel.uah.es/en/lab/cybertory/analysis/trans.htm) to identify the stable ARE motifs of ERBB2 3’UTR (**Fig. S3**). 3’UTR cDNA sequencing data deposited (10.5281/zenodo.6968947).

#### Design of stable destabilized 3’UTR ARE of ERBB2

To design the destabilized 3’UTR ARE of ERBB2, we replaced the consensus stabilized UUUUU, UUGCAUGG, CCUUACAC ARE motifs in the mRNA with destabilized ARE consensus motifs of CCUC and CUGC, UAAGUUAU (**Fig. S4**).

Next, we modified some residues on the 3’ end of the 3’UTR to increase stability. Torabi S.F.et al Science 2021 (***26***), used structural biology and biochemical assays to delineate residues of helices of the nucleic acids on the PolyA tail of 3’UTR mRNA that determines their strong, moderate stability as well as rapid degradation. With this information, we analyzed the loop structures of the stabilized and destabilized 3’UTR ARE of ERBB2 (***27***) (**Fig. S5**). The stabilized ERBB2 ARE structures have stability determining U-A on the lower loop stem as well as on the lower URIL. Using the mutated residues from the (***26***), that confers moderate degree stability as an example. In designing the destabilizing ERBB2 3’UTR (we changed the M5 (C-U), M6 (U-C) and M11 (A-U) all residues marked in red asterisks). Changes in these residues have a remarkable impact on the stability of transcript up to 120mins (***26***). Having estabilished the structure of stabilized ERBB2 3’UTR and mutated residue of destabilized ERBB2 3’UTR that increased their stability. We incoporated into these destabilized constructs with an upstream 5’ BstB1 restriction site followed by a polyA sequence to stop the RFP transcription of the vector and then followed by a DCP1A promoter. At the 3’ end we added a polyA sequence followed by BamH1 restriction site (**Fig. S6**).

#### Synthesis of destabilized 3’UTR ARE of ERBB2 as gblock

We synthesized the destabilized 3’UTR ARE of ERBB2 as gblock from Integrated DNA Technologies (IDT Inc USA).

#### Cloning of destabilized 3’UTR ARE of ERBB2 into Sp6 vector

To clone the destabilized 3’UTR ARE of ERBB2 into pLenti-CMVSP6-nEGFP-SV40-PURO (Addgene: # 138364).

1. **Miniprep of vector** A stab of the vector was inoculated into LB media and grown over night shaking at 250rpm at 37°C. The plasmid vector gDNA was extracted using the Qiagen Midikit (Cat no: 12943).
2. **Digest of vector with BstBI and BamH1** The vector was digested in BstBI and BamH1 at 37°C overnight using NEB buffer 2.1

1µl (400ng) of vector gDNA

1µl of Bstb1 and Bamh1 respectively

3µl of buffer

25µl of H20

1. **Gel extraction of vector**

The digested vector was gel extracted using Qiagen gel extraction kit (Cat no: 28704)

**Insertion of the gblock of the destabilized ERBB2 3’UTR ARE into the digested vector.**

1. **Digest insert with BstB1 and BamH1**

The synthetic gblock containing the destabilized ERBB2 3’UTR ARE were digested with BstB1 and BamH1 in NEB buffer 2.1 for 1hr at 37°C

1µl (200ng) of gblock DNA

1µl of Bstb1 and Bamh1 respectively

3µl of buffer

25µl of H20

1. **Gel extraction of insert**

The digested synthetic gblock containing the destabilized ERBB2 3’UTR was gel extracted using Qiagen gel extraction kit (Cat no: 28704)

#### Ligation

2µl T4 ligase buffer

0.5µl Plasmid vector

4µl Insert (digested synthetic gblock)

20µl Nuclease free water

1µl T4 ligase (NEB)

At 22°C for 3hrs

#### Transformation

We transformed competent recombination efficient, E-coli (NEB 5 alpha, Cat no: C298H) using SOC media. 25µl of each cell was incubated on ice with 5µl of the ligation mix for 30mins. After 30mins was heat shocked in water bath at 42°C for 30sec. Tubes were placed back on ice for 2mins and 900µl of SOC media was added and then incubated at 37°C shaking for 1hr at 250rpm. After which they were plated on LB Agar plate containing ampicillin and the plate was incubated overnight at 37°C

#### Colony picking and Miniprep

Colonies were picked with pipette tips and inoculated into 5ml LB media containing Ampicillin and grown overnight at 37°C shaking at 250rpm. The pellets were spurned down and gDNA was extracted using the Qiagen Midikit (Cat no: 12943). And using a nanodrop machine the gDNA were quantified.

#### Colony PCR with ERBB2 3’UTR primers and DCP1A promoter primers

We performed PCR with ERBB2 primers and DCP1A primers using the PCR cycle described above.

#### Gel extraction

The colony PCR products of ERBB2 3’UTR and DCP1a promoters was gel extracted using Qiagen gel extraction kit (Cat no: 28704)

#### Sanger sequencing

We sequenced the cloned synthetic gblock amplicons of ERBB2 3’UTR using the primers described above by Sanger Sequencing with Psomagen Inc, Brooklyn New York. By cloning we obtained 4 clones that matched to the ERBB2 3’UTR: des (destabilized) ARE ERBB2-1, des ARE 3’UTR ERBB2-2, des ARE 3’UTR ERBB2-3, des ARE 3’UTR ERBB2-4 (**Fig. S7**)

#### Design of Gibson Assembly Primers

##### Gibson Assembly

We succeeded in cloning the destabilized ERBB2 3’UTR into the vector by ligation, however, to extend this approach, we used GIBSON Assembly and succeeded in cloning ERBB2 destabilized 3’UTR synthetic constructs into the plasmid vector.

#### ERBB2 and Vector Gibson Assembly primers design

We loaded the sequences of the synthetic gblock of destabilized ARE 3’UTR ERBB2 and vector (Sp6 vector) into the NEB builder (https://nebuilder.neb.com/<u>#!/</u>) which generated the following primers to amplify the ERBB2 and Sp6 vector primers for Gibson Assembly

Sp6 vector forward: ggccccgccgTAATGTGAGTTAGCTCACTCATTAGG

Sp6 vector Reverse: cctcgcgggcATTGCGTTGCGCTCACTG

ERBB2 Forward: gcaacgcaatGCCCGCGAGGACCCGCCC

ERBB2 Reverse: actcacattaCGGCGGGGCCGGCCCCTA

#### Gibson Assembly of the destabilized 3’UTR ARE of ERBB2

##### PCR amplification Vector (Sp6) and Insert (destabilized ARE 3’UTR ERBB2)

6µl H20

1µl of ERBB2/Sp6 vector Forward primers respectively into a tube

1µl of ERBB2/Sp6 vector Reverse primers respectively into a tube

2µl of gDNA (200ng).

10µl of Q5 high fidelity polymerase 2X master mix (NEB cat: M0492S).

##### Gibson Assembly PCR cycle

98°C for 30sec

98°C for 10sec

55°C for 30sec

68-72 for 1min

Step2-3 for 38 cycles

4°C hold

##### NEB HIFI DNA Assembly

0.5µl of Sp6 vector mixed with 4µl of insert (destabilized ARE 3’UTR ERBB2).

10µl of NEB HiFi DNA assembly master mix (NEB cat no: E2621S).

5.5µl of deionized water

Placed at 50°C for 15mins.

#### Transformation Recombination deficient E. coli (NEB Cat no C3019H)

We transformed deficient competent E-coli with the DNA assembly using SOC media 25µl of each cell was incubated on ice with 5µl of the ligation mix for 30mins. After 30mins was heat shocked in water bath at 42°C for 30sec. Tubes were placed back on ice for 2mins and 900ul of SOC media was added and then incubated at 37°C shaking for 1hr at 250rpm. After which they were plated on LB Agar plate containing ampicillin and the plate was incubated overnight at 37°C

#### Colony picking and Miniprep

Colonies were picked with pipette tips and inoculated into 5ml LB media containing Ampicillin and grown overnight at 37°C shaking at 250rpm. The pellets were spurned down and gDNA was extracted using the Qiagen Midikit (Cat no: 12943). And using a nanodrop machine the gDNA were quantified.

#### Colony PCR with ERBB2 Gibson primers

We performed PCR with ERBB2 Gibson primers by using the PCR cycle described above.

#### Gel extraction

The colony PCR products of ERBB2 3’UTR Gibson assembled was gel extracted using Qiagen gel extraction kit (Cat no: 28704)

#### Sanger sequencing of the cloned destabilized 3’UTR ARE of ERBB2 amplicon

We sequenced colony PCR product of Gibson assembled ERBB2 3’UTR using the Gibson primers described above by Sanger Sequencing with Psomagen Inc, Brooklyn New York. We succeed in a obtaining a clone desARE3’UTR ERBB2-30 (**Fig. S8**).

#### Sequence alignment of wildtype cDNA, RNA versus the destabilized ARE 3’UTR ERBB2 cDNA and RNA sequences

To confirm the engineered changes of the destabilized ARE 3’UTR of ERBB2 versus the wildtype ERBB2 cDNA and RNA, we aligned the wildtype and the engineered destabilized and cloned sequences both on the RNA level with their control wildtype using the software Clustalw Omega (https://www.ebi.ac.uk/Tools/msa/clustalo/) (**Fig. S4**).

#### Transfection/ electroporation of destabilized 3’UTR ARE of ERBB2 into BT474 wildtype, BT474 clone 5 (Trastuzumab resistant) and NCI H1975, NCI-H2030, and HCT116 cancer cells

To introduce the destabilized 3’UTR into the cancer cells, we electroporated constructs into 200,000 to 500,000 cells with 50ng of plasmid containing the constructs using the BioRad electroporation system. We used the preset mammalian protocol set for 293T cells and pulsed the cells in a cuvette 2x. Cells were then seeded into 6 well plates and viewed for morphology and red fluorescent protein (RFP) expression in 24hrs. After 24hrs the cells expressed RFP indicating that the constructs were successfully integrated into the cells.

##### YES1 overexpression

YES1 Y537F overexpression vector (Addgene: 51299) was transfected into desARE3’UTR ERBB2-3 and 30 cells that have lost ERBB2 and YES1 by introduction of destabilized desARE3’UTR ERBB2. Overexpression of YES1 was confirmed by YES1 western blot.

##### Cell Microscopy

We observed the cells regularly under the light microscope at 20x. Within day 4, the BT474 wildtype containing the destabilized elements showed distorted and ruptured membrane compared to the control. For the trastuzumab resistant cells, the changes in cellular morphology were observed starting Day 9 for the cells containing destabilized constructs. The cell size was drastically reduced compared to controls

#### Immunofluorescence

To determine the ERBB2, Caspase 3,9 and cleaved caspase 3 protein expression changes in the destabilized breast and lung cancer compared to the wildtype. We seeded the cells on a tissue culture slide or 6 well plate at 2,000-3,000 per well. The cells were allowed to grow for 1 day and after they were fixed with 200µl of 4% paraformaldehyde added to the cell media. The cells were placed at 4°C for 5mins, after which the paraformaldehyde was decanted. Anti-ERBB2 (Human ErbB2/Her2 Mab, Clone 191924 R&D systems) 1:1000 in BSA were added to the slide wells and sealed with aluminum foil and kept at 4°C overnight. After which the antibody is removed and secondary antibody Alexa 488 (green) or Alexa 610 (red) were added at 1:10,000 and covered with aluminum foil and kept at room temperature for 1hr. After 1hr, the secondary dye was washed with PBS (phosphate buffer saline) 3x and then with a Kim wipes the edges were wiped off water. And then a drop of DAPI (Nuceloblue -nuclear stain) was added and then sealed with cover slips and the slides were viewed under a Nikon confocal microscope. For the tissue slides, slides returned from the expert pathologists were washed in 1x PBS by dipping and subsequently, dripped off and cleaned with Kim wipes. Anti-ERBB2, CNOT1, XRN1 and UPF3B (1:1000) were dropped on the slides and covered overnight in 4°C to avoid desiccation. Slides were washed in 1x PBS and Alexa 610 were added and incubated at room temp for 1hr, after which the slides were washed and a drop of DAPI added and covered with a coverslip and image were acquired under a Nikon confocal microscope.

#### Western blot

To quantify the ERBB2, YES1, WNK1 and CNOT1 protein expression changes in wildtype cells compared to the cells containing the destabilized 3’UTR ARE. We harvested the cells and lysed them in cocktail of protease inhibitor in MPER buffer. The protein extract was stored in -80C until use. The protein was separated in 12% stacked SDS page gels and separated by initial run at 75 volts and after 15mins at 120 V until complete separation. The proteins were transferred unto a PVDF membrane already charged with methanol at 25 V for 1hr. Membrane was blocked with BSA and subsequently incubated with anti-ERBB2(Human ErbB2/Her2 MAb Clone 191924) or anti-YES1 and WNK1 (Cell signaling #65890 and #4979), CNOT1 (14276-1-AP) overnight in BSA. Primary antibody was removed and secondary anti-mouse IR 800CW dye (Li-COR) was added at 1:1000 incubated with BSA covered from light for 2hrs. After incubation, membrane was washed with TBST 3x and image was taken on Li-COR Oddysey chemilumiscent imager.

#### Cell Viability

To quantify if the destabilized constructs affect the cell survival. We performed cell viability comparing the wildtype cells with the cells carrying destabilized constructs with cell titre glo (Promega cat: G7570).

#### Caspase 3/7 assay, caspase 3, cleaved caspase 3 and caspase 9 protein analysis

To assay for increased caspase activity, we used the caspase 3/7 glo kit (Promega cat: G8090) as well anti-Caspase 3, 9 (Abcam #4051, #25758), cleaved caspase 3 (Cell Signaling #9664). The kit was used according to the manufacturer’s instructions. We performed immunofluorescence with antibodies against cleaved caspase 3 and caspase 9 on wildtype cells and the engineered destabilized cells as described above.

#### Phosphorylation kinase array

To determine the how the kinases are affected by the destabilization of the ERBB2 protein. We used the human phospho-kinase array kit (Cat no: ARY003C). We followed the protocol as described.

#### Migration assay

Wounds were made on the cell monolayer using a sterile 200 μL pipette tip. Cells were washed with 1× PBS and fresh media were added to each well following the wound. Images of scratched areas were taken at 10× magnification using an AE30 inverted microscope (Motic, Richmond, BC, Canada).

#### RNA Sequencing

To confirm the loss of ERBB2 on genome scale following the destabilization of the 3’UTR ARE as well as to ascertain to what level the ERBB2 interactome is affected by loss of ERBB2 and it’s specificity compared to the wild type controls. We performed RNA Seq on the RNA from desARE 3’UTR ERBB2-3 and desARE 3’UTR ERBB2-30 compared to the wildtype and vector controls. RNA Seq analysis was performed using established pipelines and Biojupies (https://maayanlab.cloud/biojupies/). Data deposited: 10.5281/zenodo.6968947

#### Quantitative Reverse Transcriptase PCR (qPCR)

To quantify the ERBB2 expression changes upon destabilization of the ERBB2 3’UTR ARE comparing the wildtype and the vector controls. We did quantitative PCR on ERBB2, deadenylases CNOT1, XRN1 and PARN as well as on decapping enzyme DCP1A using their exon primers and GAPDH as control. All primers used are listed **List S1.**

##### Optical genome mapping

To understand if our engineering method introduced copy number variation, indel etc. We used optical genome mapping provided by Bionano.

(https://bionanogenomics.com/products/saphyr/). The experiments were performed in two independent replicates. Data deposited :10.5281/zenodo.6968947

#### Animal Study

We obtained IACUC approval of the animal study from institutional board. 25 female NSG mice were purchased from the Jackson laboratory. Animals were received and allowed to acclimate according to institutional protocol. NCI H1975 were expanded as described above and on the day of implantation, cells were harvested, washed, and resuspended in phosphate buffer saline (PBS). The confluency of the cells was 80%. We implanted 5million cells in PBS on the flank of each mouse. After 35 days, huge tumors engrafted and on 36 days, we randomize the mice into 5mice per cage into 5 groups and we ensured equal distribution of tumor size within each group. The vector and desARE3’UTRERBB2-1, 3 and 30 were administered at 20µg per mice intraperitoneally 12hrly for 9days with a one-day dosing break. The group of wildtype NCI H1975 mice received no treatment. Daily measurement of tumor size (length and width), weight and body condition score were obtained. At day 46, due to the enormous size of the wildtype and vector tumor sizes, the control groups were euthanized and subsequently the mice receiving the constructs were also euthanized to perform complete necropsy, full blood count and blood chemistry analysis. The necropsy and complete blood count and histopathology were performed by expert pathologist from the Memorial Sloan Kettering Hospital, New York. They were blinded to the experimental details

#### Full blood count and electrolyte analysis

The full blood count and necropsy and electrolyte analysis were performed by expert pathologist at the Memorial Sloan Kettering Cancer Center, New York City. See **List S4, 5**.

#### Image J

We used the software Image J to quantify the immunofluorescence signal of ERBB2, Cleaved Caspase 3 and Caspase 9

#### GraphPad Prism

We used the GraphPad Prism software to draw all the bar charts presented and performed all statistics with the software.

## Notes

### Competing Interest Statement

Chidiebere U Awah, Kevin Struhl and Olorunseun O Ogunwobi have filed for a patent based on findings from this work. Chidiebere U Awah, Kevin Struhl, Dan Weiser, and Olorunseun O Ogunwobi are Co-Founders of UTR Therapeutics Inc, a start-up company with interest in this work.

## References

1. https://www.who.int/news-room/fact-sheets/detail/cancer

2. RL Siegel, KD Miller, A Jemal, Cancer statistics 2020, CA Cancer J Clin 70, 7–30 (2020).

3. H Sung, J Ferlay, RL Seigel, M Laversanne, I Soerjomatarm, A Jemal, F Bray, Global cancer statistics 2020: Globocan estimates of incidence and mortality worldwide for 36 cancers in 185 countries. CA Cancer J Clin 71, 209–249 (2021).

4. F Zhao, B Copley, Q Niu, Liu Q, F Liu, J A Johnson, T Sutton, G Khramtsova, E Sveen, TF Yohismatsu, Y Zheng, A Ibraheem, N Jaskowiak, R Nanda, G F Fleming, OI Olopade, D Huo: Racial disparities in survival outcomes among breast cancer patients by molecular subtypes. Breast Cancer Res Treat 185, 841–849 (2020).

5. D J Slamon, G M Clark, SG Wong, WJ Levin, A Ullrich, W L McGuire, Human breast cancer: correlation of relapse and survival with amplification HER-2/neu oncogene. Science 235, 177–82 (1987).

6. Hynes NE, Lane HA: ERBB2 receptors and cancers: the complexity of targeted inhibitors. Nat.Rev Cancer 5, 341–354 (2005).

7. M A Molina, J Codony-Servat, J Albanell, F Rojo, J Arribas, J Baselga. Trastuzumab (herceptin), a humanized anti-ERBB2 receptor monoclonal antibody, inhibits basal and activated ERBB2 ectodomain cleavage in breast cancer cells. Cancer Res 12, 4744–9 (2005)

8. M A Cobleigh, C L Vogel, D Tripathy, N J Robert, S Scholl, L Fehrenbacher, J M Wolter, V Paton, S Shak, G Lieberman, D J Slamon. Multinational study of the efficacy and safety of humanized anti-ERBB2 monoclonal antibody in women who have ERBB2-overexpressing metastatic breast cancer that has progressed after chemotherapy for metastatic disease. J Clin Oncol 9, 2639–48 (1999).

9. Y Zhang. The root cause of drug resistance in ERBB2-positive breast cancer and the therapeutic approaches to overcoming the resistance. Pharmacol Ther 218, 107677 (2021).

10. R Bose, C X Ma. Breast Cancer, *ERBB2* Mutations, and Overcoming Drug Resistance. N Engl. J Med 385, 1241–1243 (2021).

11. AB Hanker, BP Brown, J Meiler, A Marín, H S Jayanthan, D Ye, C-C Lin, H Akamatsu, K-M Lee, S Chatterjee, D R Sudhan, A Servetto, M R Brewer, J P Koch, J H Sheehan, J He, A S Lalani, C L Arteaga. Co-occurring gain-of-function mutations in ERBB2 and HER3 modulate ERBB2/HER3 activation, oncogenesis, and ERBB2 inhibitor sensitivity. Cancer Cell 39,1099–1114 (2021).

12. A Derakhshani, Z Rezaei, H Safarpour, M Sabri, A Mir, M A Sanati, F Vahidian, Ali G Moghadam, A Aghadoukht, K Hajiasgharzadeh, B Baradaran. Overcoming trastuzumab resistance in ERBB2-positive breast cancer using combination therapy. J Cell Physiol 4, 3142–3156 (2020).

13. B T Li, E F Smit, Y Goto, K Nakagawa, H Udagawa, J Mazières, M Nagasaka, L Bazhenova, A N Saltos, E Felip, J M Pacheco, M Pérol, L P-Ares, K Saxena, R Shiga, Y Cheng, S Acharyya, P Vitazka, J Shahidi, D Planchard, P A Jänne, DESTINY-Lung01 Trial Investigators. Trastuzumab Deruxtecan in ERBB2-Mutant Non-Small-Cell Lung Cancer. N Engl J Med 386, 241–251 (2022).

14. H P Safran, K Winter, D H Ilson, D Wigle, T DiPetrillo, M G Haddock, T S Hong, L P Leichman, L Rajdev, M Resnick, L A Kachnic, S Seaward, H Mamon, D A D Pardo, C M Anderson, X Shen, A K Sharma, A W Katz, J Salo, K L Leonard, J Moughan, C H Crane. Trastuzumab with trimodality treatment for oesophageal adenocarcinoma with ERBB2 overexpression (NRG Oncology/RTOG 1010): a multicentre, randomised, phase 3 trial. Lancet Oncol 23, 259–269 (2022).

15. M Takeda, K Nakagawa. First-and Second-Generation EGFR-TKIs Are All Replaced to Osimertinib in Chemo-Naive *EGFR* Mutation-Positive Non-Small Cell Lung Cancer? Int J Mol Sci 20, 146 (2019).

16. A J, P-Vallillo, LV Sequist, Z Piotrowska. Emerging Treatment Paradigms for EGFR-Mutant Lung Cancers Progressing on Osimertinib: A Review. J Clin Oncol 38, 2926–2936 (2020).

17. J-C Soria, Y Ohe, J Vansteenkiste, T Reungwetwattana, B Chewaskulyong, K H Lee, A Dechaphunkul, F Imamura, N Nogami, T Kurata, I Okamoto, C Zhou, B C Cho, Y Cheng, E K Cho, P J Voon, D Planchard, W-C Su, J E Gray, S-M Lee, R Hodge, M Marotti, Y Khazanov, S S Ramalingam, FLAURA Investigators. Osimertinib in Untreated EGFR-Mutated Advanced Non-Small-Cell Lung Cancer. N Engl J Med 378, 113–125 (2018).

18. H Sun, Y Wu. Osimertinib in first line setting: preventive or delayed T790M occurrence? Trans Lung Cancer Res 7, S187–190 (2018).

19. S L Monica, D Cretella, M Bonelli, C Fumarola, A Cavazzoni, G Digiacomo, L Flammini, E Barocelli, R Minari, N Naldi, P G Petronini, M Tiseo, R Alfieri. Trastuzumab emtansine delays and overcomes resistance to the third-generation EGFR-TKI osimertinib in NSCLC EGFR mutated cell lines. J Exp Clin Cancer Res 36, 174 (2017).

20. C Zhu, W Zhuang, L Chen, W Yang, W-B Ou. Frontiers of ctDNA, targeted therapies, and immunotherapy in non-small-cell lung cancer. Transl. Lung Cancer Res 9, 111–138 (2020)

21. ZH Tang, JJ Lu Osimertinib resistance in non-small cell lung cancer: Mechanisms and therapeutic strategies. Cancer Lett 420, 242–246 (2018).

22. C E Vejnar, M Messih, C M Takacs, V Yartseva, P Oikonomou, R Christiano, M Stoeckius, S Lau, M T Lee, J-D Beaudoin, D Musaev, H D-Codore, T C Walther, S Tavazoie, D Cifuentes, A J Giraldez. Genome wide analysis of 3’ UTR sequence elements and proteins regulating mRNA stability during maternal-to-zygotic transition in zebrafish. Genome Res 7, 1100–1114 (2019).

23. R Geissler, A Simkin, D Floss, R Patel, E A Fogarty, J Scheller, A Grimson. A widespread sequence-specific mRNA decay pathway mediated by hnRNPs A1 and A2/B1. Genes Dev 30, 1070–85 (2016).

24. J V Geisberg, Z Moqtaderi, X Fan, F Ozsolak, K Struhl. Global analysis of mRNA isoform half-lives reveals stabilizing and destabilizing elements in yeast. Cell 156, 812–24 (2014).

25. J Lykke-Andersen, E Wagner. Recruitment and activation of mRNA decay enzymes by two ARE-mediated decay activation domains in the proteins TTP and BRF-1. Genes Dev. 19, 351–361 (2005).

26. S F Torabi, A T Vaidya, K T, Tycowski, S J DeGregorio, J Wang, MD Shu, TA Steitz, J A Steitz. RNA stabilization by a poly(A) tail 3’-end binding pocket and other modes of poly(A)-RNA interaction. Science 371, 6523 (2021).

27. J S Reuter, D H Matthews. RNAstructure: software for RNA secondary structure prediction and analysis. BMC Bioinformatics 11, 129 (2010).

28. T Kute, C M Lack, M Willingham, B Bishwokama, H Williams, K Barrett, T Mitchell, JP Vaughn. Development of Herceptin resistance in breast cancer cells. Cytometry A 57, 86–93 (2004).

29. S Zazo, P G-Alonso, E M-Aparicio, C Chamizo, I Cristóbal, O Arpí, A Rovira, J Albanell, P Eroles, A Lluch, J M-Gúrpide, F Rojo. Generation, characterization, and maintenance of trastuzumab-resistant ERBB2+ breast cancer cell lines. Am J Cancer Res 6, 2661–2678 (2016).

30. L Wang, Q Wang, P Xu, L Fu, Y Li, H Fu, H Quan, L Lou. YES1 amplification confers trastuzumab-emtansine (T-DM1) resistance in ERBB2-positive cancer. Br J Cancer 123, 1000–1011 (2020).

31. A B Jaykumar, J U Jung, P K Parida, T T Dang, C Wichaidit, A R Kannangara, S Earnest, E J Goldsmith, G W Pearson, S Malladi, M H Cobb. WNK1 Enhances Migration and Invasion in Breast Cancer Models. Mol. Cancer. Ther 20, 1800–1808 (2021).

32. J Rosenbluh, D Nijhawan, A G Cox, X Li, J T Neal, E J Schafer, T I Zack, X Wang, A Tsherniak, A C Schinzel, D D Shao, S E Schumacher, B A Weir, F Vazquez, G S Cowley, D E Root, J P Mesirov, R Beroukhim, C J Kuo, W Goessling, W C Hahn. β-Catenin-driven cancers require a YAP1 transcriptional complex for survival and tumorigenesis. Cell 151, 1457–73 (2012).

33. FJ Vizeacoumar, R Arnold, FS Vizeacoumar, M Chandrashekhar, A Buzina, J T F Young, J H M Kwan, A Sayad, P Mero, S Lawo, H Tanaka, K R Brown, A Baryshnikova, A B Mak, Y Fedyshyn, Y Wang, G C Brito, D Kasimer, T Makhnevych, T Ketela, A Datti, M Babu, A Emili, L Pelletier, J Wrana, Z Wainberg, P M Kim, R Rottapel, C A O’Brien, B Andrews, C Boone, J Moffat. A negative genetic interaction map in isogenic cancer cell lines reveals cancer cell vulnerabilities.Mol Syst Biol 8, 696 (2013).

34. T Hart, M Chandrashekhar, Aregger M, Z Steinhart, KR Brown, MacLeod G, M Mis, M Zimmermann, A Fradet-Turcotte, S Sun, P Mero, P Dirks, S Sidhu, FP Roth, OS Rissland, D Durocher, S Angers, J Moffat. High-Resolution CRISPR Screens Reveal Fitness Genes and Genotype-Specific Cancer Liabilities. Cell 163, 1515–26 (2015).

35. T D Martin, D R Cook, M Y Choi, M Z Li, K M Haigis, S J Elledge. A Role for Mitochondrial Translation in Promotion of Viability in K-Ras Mutant Cells. Cell Rep 20,427–438 (2017).

36. J L Lühmann, M Stelter, M Wolter, J Kater, J Lentes, A K Bergmann, M Schieck, G Göhring, A Möricke, G Cario, M Žaliová, M Schrappe, B Schlegelberger, M Stanulla, D Steinemann. The Clinical Utility of Optical Genome Mapping for the Assessment of Genomic Aberrations in Acute Lymphoblastic Leukemia. Cancers 17, 4388 (2021).

37. XH Zhang, LY Tee, XG Wang, QS Huang, SH Yang. Off target Effects in CRISPR/Cas9-mediated Genome Engineering. Mol Ther Nucleic Acids 11, e264 (2015)

38. A Fire, S Xu, M K Montgomery, S A Kostas, S E Driver, C C Mello. Potent and specific genetic interference by double-stranded RNA in Caenorhabditis elegans. Nature 391,806–11 (1998).

39. R J Dohmen, P Wu, A Varshavsky. Heat-inducible degron: a method for constructing temperature-sensitive mutants. Science 263, 1273–6 (1994).

40. K M Sakamoto, K B Kim, A Kumagai, F Mercurio, C M Crews, R J Deshaies. Protacs: chimeric molecules that target proteins to the Skp1-Cullin-F box complex for ubiquitination and degradation. Proc Natl Acad Sci U S A 98, 8554–9 (2001).

41. A Yesbolatova, Y Saito, N Kitamoto, H M-Itou, R Ajima, R Nakano, H Nakaoka, Kosuke Fukui, K Gamo, Y Tominari, H Takeuchi, Y Saga, K-I Hayashi, M T Kanemaki. The auxin-inducible degron 2 technology provides sharp degradation control in yeast, mammalian cells, and mice. Nat Commun 11, 570 (2020).

